# A Global Analysis of Mutations Accompanying Microevolution in the Heterozygous Diploid Pathogen *Candida albicans*

**DOI:** 10.1101/288597

**Authors:** Iuliana V. Ene, Rhys A. Farrer, Matthew P. Hirakawa, Kennedy Agwamba, Christina A. Cuomo, Richard J. Bennett

## Abstract

*Candida albicans* is a heterozygous diploid yeast that is a commensal of the human gastrointestinal (GI) tract and a prevalent opportunistic pathogen. Here, whole-genome sequencing was performed on multiple *C. albicans* isolates passaged in different niches to characterize the complete spectrum of mutations arising during microevolution. We reveal that evolution during short time-scales (<600 generations) is driven by both *de novo* base substitutions and short-tract loss of heterozygosity (LOH) events. In contrast, large-scale chromosomal changes are relatively rare, although chromosome 7 trisomies repeatedly emerged during passaging in one GI colonization model. Both strain background and chromosomal features affected mutational patterns, with mutation rates being greatly elevated in regions adjacent to emergent LOH tracts. Mutation rates were also elevated during host infection where genomes showed strong evidence of purifying selection. These results establish the genetic events driving *C. albicans* evolution and that this heterozygous diploid is extensively shaped by purifying selection.

## Introduction

Microbial evolution studies provide insights into genome dynamics and the factors that shape genome evolution (1–5). Most studies have focused on haploid or homozygous diploid genomes, yet there is an increasing interest in defining genome variation in heterozygous diploid genomes (4, 6–9). Sexual reproduction often plays an important role in accelerating adaptation in eukaryotic species, as meiotic recombination promotes genome rearrangements and mutation rates are elevated compared to vegetative cells (5, 10, 11). However, mitotic populations can still generate diversity by a number of mechanisms including *de novo* base substitutions, loss of heterozygosity (LOH) events, and insertion or deletions (indels), as well as larger rearrangements and even the acquisition of chromosomal aneuploidies. Many of these events have been defined in model eukaryotes and are also recognized as major drivers of somatic mosaicism and cancer development in the human genome (12, 13).

In this study, we provide a high-resolution picture of genome microevolution in the heterozygous diploid yeast *Candida albicans*. *C. albicans* is an opportunistic pathogen responsible for a variety of debilitating mucosal infections and life-threatening systemic infections (14, 15). A common resident of the human microbiota, *C. albicans* can become pathogenic in individuals that are immunocompromised or undergo prolonged antibiotic use (15–17). The genome consists of 8 chromosomes with the reference isolate SC5314 containing ~70,000 heterozygous positions representing ~0.5% of the 14.3 Mb genome (18–20). Heterozygous regions of the genome can undergo LOH which is increased in response to stressful environments (21–24). Isolates can also experience large-scale changes including the acquisition of aneuploid forms (24–27). *C. albicans* isolates therefore display extensive genomic plasticity due to a variety of events including acquisition of SNPs and indels, LOH events, and changes in gene and chromosome copy number (22, 24, 27–32).

Here, we define the complete spectrum of mutations accompanying microevolution in *C. albicans*. We performed deep sequencing on multiple clinical isolates to precisely determine the mutations arising during both *in vitro* and *in vivo* passaging. Our experiments reveal that microevolution is driven by widespread, small-scale genetic changes, overwhelmingly represented by *de novo* base substitutions and short-tract LOH events. In contrast, large-scale genomic changes are rare, although both long-tract LOH events and the acquisition of supernumerary chromosomes were observed, with the latter found to be a niche-specific alteration. We also identify hypermutable domains within the genome including repetitive and telomeric regions. Furthermore, we show that DNA recombination events are themselves highly mutagenic and contribute to genomic variation by introducing a large number of *de novo* mutations. Genetic events leading to gains and losses of heterozygosity occurred at similar rates so that global heterozygosity levels were, in most cases, stably maintained throughout microevolution. Finally, we demonstrate that mutational patterns reveal a dominant role for purifying selection, with emergent mutations that alter protein-coding sequences often purged from the genome during infection of the mammalian host.

## Results

### Microevolution of *C. albicans* diploid genomes

We selected four clinical isolates of *C. albicans* (SC5314, P78048, P76055 and P57055) for microevolution experiments. These isolates belong to three major *C. albicans* clades (I, I, II and III, respectively), exhibit normal fitness and morphology, and have heterozygous diploid genomes (with no chromosomal aneuploidies). Heterozygous positions represent 0.41% to 0.55% of these genomes (Supplementary Tables 1-3) (24). Strains were passaged both *in vitro* and in three different murine models *in vivo* (Figure 1A). The latter included two commensal models of gastrointestinal (GI) colonization using either a standard diet (SD) that requires antibiotics for *C. albicans* colonization (33) or a purified diet (PD) that does not require antibiotics for stable colonization (34). A model of systemic infection was also utilized in which fungal cells were introduced into the murine tail vein and subsequently recovered from the kidney, the major organ targeted by *C. albicans* (35). For GI colonization, fungal cells were collected from fecal pellets after 42 days (~227 generations, *n*=2-3). For systemic infection, five sequential passages were performed in which fungal cells were isolated from infected kidneys three days post infection and used for infection of new hosts (~240 generations, *n*=2). For comparison, *in vitro* passaging was performed daily under standard laboratory conditions (YPD medium, 30°C) and isolates collected after 80 days (~600 generations, *n*=1-2). The genomes of evolved isolates were analyzed by Illumina ultra-deep sequencing (average of 185X coverage, 97.7% of SC5314 assembly covered by reads, see Supplementary Table 2 and Materials and Methods). Using high depth read alignments and stringent variant calling with Haplotype Caller (GATK) and Pilon (36), single nucleotide polymorphisms (SNPs), heterozygous positions and indels were identified for each isolate (Supplementary Table 2), and a number of these were further validated as described below.

**Figure 1.**
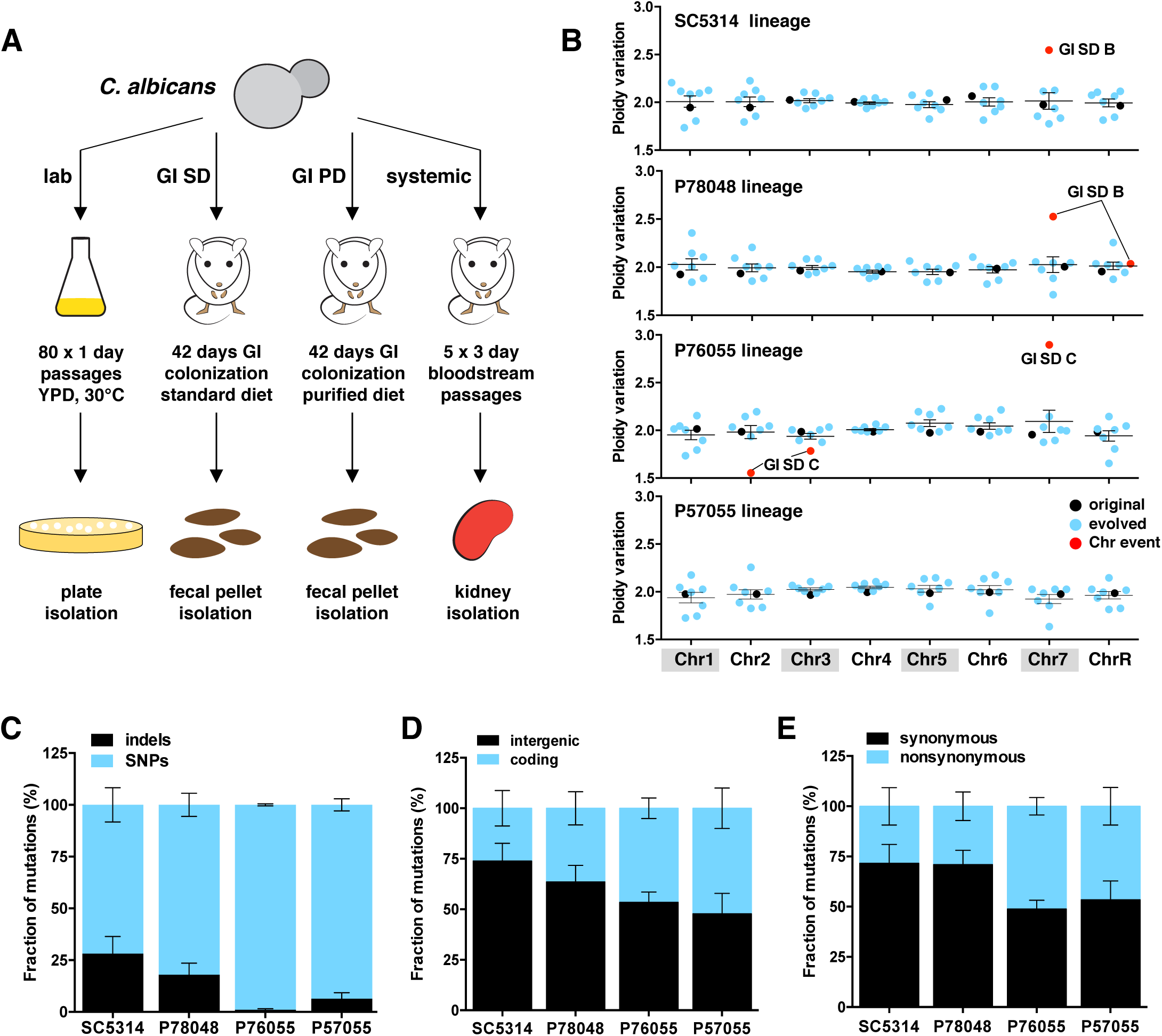
Microevolution of *C. albicans* genomes. (A) Schematic of *in vitro* and *in vivo* microevolution experiments. GI SD, gastointestinal standard diet; GI PD, gastointestinal purified diet. (B) Ploidy variation based on read depth for each evolved isolate and each chromosome, normalized by the average genomic read depth. Isolates with significant ploidy changes (including full and segmental aneuploidies) as well as those with large chromosomal events are marked in red. (C) Distribution of SNPs and indels, intergenic and coding mutations, synonymous and nonsynonymous mutations across microevolved isolates averaged for each lineage. Note that panels include both GOH and LOH events.

### Large-scale chromosomal changes acquired during microevolution

Sequence read depth across each of the 28 microevolved genomes revealed that a small subset of isolates underwent changes at the chromosomal level. Aneuploidy was observed in three out of 28 evolved isolates and in each case involved chromosome (Chr) 7 trisomies (Figure 1B and Supplementary Figure 1A). These aneuploid forms emerged in three different strain backgrounds that were each passaged in the GI SD model, suggesting a fitness benefit may be associated with Chr 7 trisomy under these growth conditions. One of the three aneuploid isolates also became monosomic for two terminal regions involving the right arm of Chr 2 (1.15 Mbp region) and the left arm of Chr 3 (0.27 Mbp region) (P76055 GI SD isolate C, Figure 1B and Supplementary Figure 1B). Previous studies have similarly observed aneuploid forms among natural isolates and that chromosome-level changes can arise during passaging or in response to antifungal treatment (24, 27, 29, 31, 32, 37, 38). An analysis of copy number variation across smaller genomic regions (100-1000 bp windows) is included in Supplementary Material and Supplementary Figures 7-8.

### Common patterns of microvariation in *C. albicans* genomes

A detailed analysis of microevolved isolates was performed to determine the spectrum of nucleotide changes in each genome. A total of 564 mutations were identified across the 28 microevolved isolates in the four lineages. All 564 mutations were individually evaluated using IGV (39) and 63 sites were additionally verified using an allele-specific fluorescent PCR technology (KASP genotyping, LGC). Of these, 55 positions (87%) matched the genotypes from genome sequencing (Supplementary Table 4). Validated mutations included both SNPs and indels in both genic and intergenic regions (Supplementary Figure 1C).

Mutations were subdivided into those leading to gains or losses of heterozygosity (GOH and LOH, respectively). GOH and LOH were further classified as resulting from either indels or changes in SNPs. For example, *in vitro* passaging of *C. albicans* isolates for 600 generations revealed a total of 31 mutations comprising 6 indels (4 insertions and 2 deletions) and 25 SNPs (17 transitions and 8 transversions) across four isolates. These 31 mutations were the result of 8 GOH events (7 *de novo* base substitutions and 1 indel) and 11 LOH events due to recombination. 19 of the mutations occurred in intergenic regions and 12 occurred in coding sequences, of which there were 7 synonymous and 5 nonsynonymous mutations.

Microevolution consistently resulted in more SNPs (both GOH and LOH events) than indels, independent of strain background or evolution niche. Thus, an average of 87.2% of mutations involved SNPs and 12.8% involved indels (Figure 1C). For GOH events, the average ratio of base substitutions to indels was 1:0.17, which is much lower than ratios reported for *Saccharomyces cerevisiae* (~1:0.03) (40–42), suggesting that *C. albicans* experiences proportionally higher rates of indels to *de novo* substitutions than *S. cerevisiae*.

An average of 41% of all mutations (including both GOH and LOH SNPs and indels) occurred in coding regions and comprised 60.3% synonymous and 39.7% nonsynonymous mutations (Figure 1D, E). Nonsynonymous mutations predicted to disrupt ORF function were rare; only 14 nonsense mutations and 5 readthrough mutations occurred across all evolved isolates, and 17 of the 19 mutations were the direct result of three very large LOH events that occurred in two microevolved lineages (described below and Supplementary Table 5). Base substitutions (GOH SNPs) in evolved isolates were the result of a higher fraction of transitions (54.4%) than transversions, with a *Ts/Tv* ratio of 1.3:1, which is lower than the 2:1 ratio reported for model yeast genomes (41, 43) (Supplementary Figure 1D). These mutational patterns were consistent across microevolved isolates revealing that they are independent of genetic background and the environment in which isolates are passaged (Figure 1C-E and Supplementary Figure 1E-G).

### Purifying selection shapes the evolution of *C. albicans* genomes

In the absence of bottlenecks, new mutations that have deleterious effects may be purged from the population via purifying selection, and we therefore tested whether mutational patterns in our dataset showed evidence for selection. If occurring randomly, mutations will accumulate in intergenic and coding regions at frequencies proportional to their representation in the genome (40, 41, 44). In our experiments, 48% of all *de novo* base substitutions (GOH SNPs) and 61.5% of GOH indels were present in intergenic regions, even though these regions account for only 36.2% of the genome (*P* < 0.05, Figure 2A). Moreover, none of the indels (0/5) found in coding sequences resulted in frameshifts (i.e., all were a multiple of 3 nucleotides), whereas only 1/8 indels observed in intergenic regions consisted of multiples of 3 nucleotides (*P* < 0.05, difference between intergenic and coding indels is significant using a binomial distribution model, Figure 2B). The fraction of synonymous to nonsynonymous mutations also differed from that expected by chance; ~25% of coding substitutions are expected to be synonymous if mutations occur randomly (40, 44) yet over 48% of base substitutions were synonymous in our dataset (*P* < 0.05, Figure 2C). This suggests that selection frequently acts to limit the accumulation of mutations that alter the protein-coding sequence.

**Figure 2.**
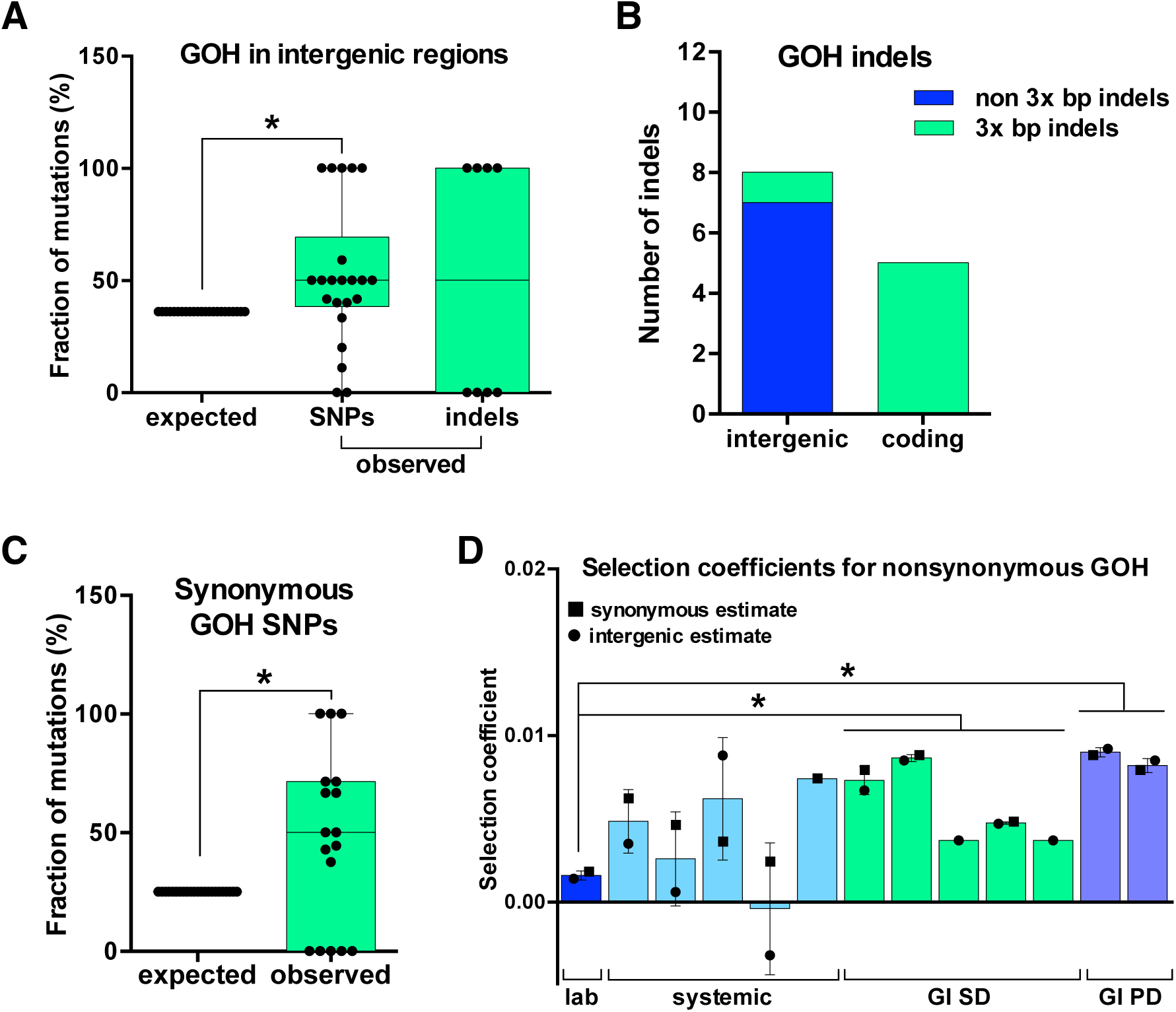
Selection shapes microevolution of *C. albicans* genomes. (A) Frequency of observed and expected GOH mutations in intergenic regions (these regions represent 36.2% of the *C. albicans* genome). GOH SNP mutations represent *de novo* base substitutions. (B) Number of GOH indels identified in coding and intergenic regions. Indels are classified based on whether they are multiple of 3 bp in length. (C) Observed and expected fractions of synonymous GOH SNPs in coding regions during microevolution experiments. ~75% of all base substitutions are expected to be nonsynonymous if they occurred randomly and were not subject to selection. (D) Selection coefficients for nonsynonymous GOH SNPs calculated based on the number of observed vs. expected nonsynonymous substitutions. Expected nonsynonymous substitutions were estimated for each microevolution experiment based on observed synonymous substitutions (squares) or observed intergenic substitutions (circles). Only isolates for which both nonsynonymous and synonymous/intergenic substitutions were observed were included in this analysis. For each panel, asterisks indicate significant differences (t-test, *P* < 0.05).

We also estimated the fraction of mutations impacted by selection by examining how many nonsynonymous mutations would be expected during microevolution based on the number of synonymous or intergenic mutations observed. Assuming an even distribution of mutational events (see Discussion), selection effectively removed an average of 71-79% of the nonsynonymous mutations predicted to occur during microevolution. Selection coefficients for nonsynonymous mutations in evolved isolates averaged 0.0053 (0.0047 using estimates based on intergenic mutations and 0.0059 using synonymous mutation rates). Selection coefficients were highest for isolates passaged in the two GI colonization models (*P* < 0.0005, Figure 2D).

Together, these results establish that *C. albicans* isolates display much higher synonymous:nonsynonymous and intergenic:genic mutation ratios than that expected by chance, implying that purifying selection removes a large fraction of the mutations impacting protein-coding genes during passaging *in vitro* and *in vivo*.

### Impact of strain background and environment on *C. albicans* mutation rates

Mutation rates were compared between the four clinical isolates and across culture conditions to examine cell-intrinsic and cell-extrinsic factors that impact microevolution. Strains passaged *in vitro* displayed an average rate of 1.17 × 10^−10^ base substitutions per base pair (bp) per generation (Supplementary Figure 2A). These *de novo* substitution rates are similar to those reported for asexual populations of *S. cerevisiae* and *Schizosaccharomyces pombe* (40–42, 45) (Supplementary Figure 2B). Mutations in *C. albicans* cells passaged *in vitro* reflected mutational patterns common to all microevolution experiments, with more frequent changes due to SNPs than to indels, and fewer mutations affecting coding regions than expected by chance (Supplementary Figure 2A). Isolates passaged *in vitro* displayed an average LOH rate of 1.61 × 10^−10^ per bp per generation, resulting from 2.75 LOH events per strain every 600 generations.

Mutation rates varied considerably depending on both the genetic background and the environment. The standard ‘laboratory’ strain SC5314 displayed the lowest mutation rates (both for GOH and LOH events) as rates in the other three lineages were 1.3 – 5.6-fold higher (Figure 3A). The environment also significantly impacted the mutation frequency; strains grown *in vivo* (either in the GI or in systemic models of infection) showed GOH rates that were 6.7 – 9.6-fold higher than those *in vitro* (Figure 3B). LOH rates were also higher *in vivo* than *in vitro* (6.8 – 12.7-fold; Figure 3B). Thus, *C. albicans* cells exhibit significantly higher mutation rates when passaged in the host (either in systemic or GI infection models) relative to *in vitro* passaging.

**Figure 3.**
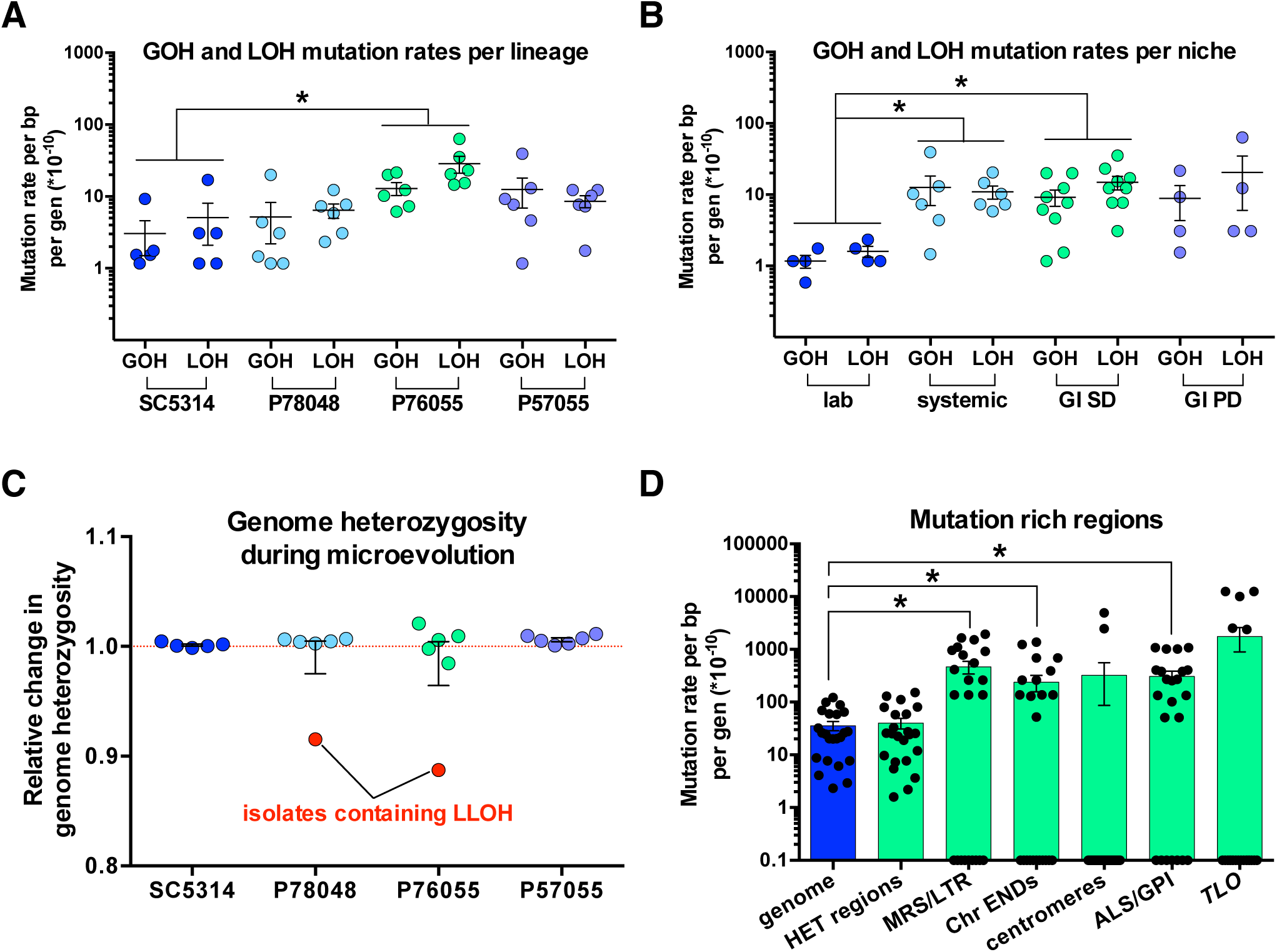
Mutation rates in *C. albicans* are impacted by strain background, environmental niche, and chromosomal features. (A,B) Effect of strain background (A) and evolution niche (B) on GOH and LOH mutation rates. Rates include both SNP and indel mutations. (C) Fluctuations in genome heterozygosity during microevolution relative to starting heterozygosity levels (red line). Major decreases in heterozygosity are only observed for isolates that underwent large-tract LOH events (LLOH; red symbols). (D) Mutation rates in specific regions of the *C. albicans* genome. These include heterozygous (HET) regions, repeat regions (MRS and LTR), Chr END regions (final 10 kb of each chromosome arm), centromeres, genes encoding *ALS* and GPI-linked proteins, and *TLO* genes. Mutations include SNPs and indels resulting from both GOH and LOH. Asterisks indicate significant differences (t-test, *P* < 0.05).

In contrast to overall GOH and LOH mutations, rates of indel formation did not differ significantly between experiments in different strain backgrounds or in different niches. The P76055 lineage displayed the lowest indel frequency (0.8 × 10^−10^ per bp per generation) compared to rates that were 3.4 - 4.4-fold higher in the other lineages (Supplementary Figure 2C). Indel rates were also elevated 3.1 – 4.6-fold in the bloodstream and in GI SD infection models relative to *in vitro* passaging, although these differences did not reach significance due to the small number of events (Supplementary Figure 2D). We note that precise *in vivo* mutation rates are difficult to determine due to approximated generation times (see Materials and Methods, Supplementary Table 3). Previous studies estimated 0.09 generations/h during systemic infection (31) and 0.14 generations/h during GI colonization (46), rates that are only 2- 3-fold lower than *in vitro* growth. It is therefore possible that these *in vivo* generation times are overestimates and mutation rates *in vivo* could be even higher than those presented here.

### Genome heterozygosity levels are maintained due to balanced GOH and LOH rates

An important question is how do *C. albicans* strains maintain genome heterozygosity levels despite frequent LOH events and the absence of conventional outcrossing (22, 24, 25, 47–49). Heterozygosity patterns were compared before and after passaging and revealed that LOH rates and GOH rates were often balanced in each experiment (Supplementary Figure 2E). Thus, genome heterozygosity levels in the four strain backgrounds were virtually unchanged following most passaging experiments, with levels within −1.5% to +2.1% of starting heterozygosity levels (Figure 3C). The exceptions to this pattern were two passaged isolates that experienced very large (>0.27 Mb) LOH events (‘LLOH tracts’) and therefore exhibited significant decreases (−8.5% and −11.3%) in heterozygous sites across their genomes. Together, these results establish that mitotic recombination events (driving LOH) and GOH events (resulting from both *de novo* base substitutions and indels) often occur at similar frequencies and that, in the absence of large LOH events, genome heterozygosity levels can be stably maintained independent of genetic background or environment.

### Defining hypervariable regions within the *C. albicans* genome

To determine the impact of genomic context on mutation rates, we compared the frequency of mutations arising at a number of chromosomal features including centromeres, terminal chromosome regions, subtelomeric *TLO* genes, *ALS* (agglutinin-like sequence) genes, other glycosylphosphatidylinositol (GPI)-linked genes, and annotated DNA repeat regions (see Supplementary Tables 6-7). Several of these features have been associated with higher mutation rates in *C. albicans* and other model organisms (4, 50–56). For each of these features, we examined the frequency of SNP and indel mutations relative to the genome average. We found that mutation rates were significantly elevated at the ends of chromosomes (6.7 fold increase within the 10 kb terminal regions), as well as in the subtelomeric *TLO* genes (48.8 fold increase) of a subset of lineages (Figure 3D and Supplementary Figure 3A,B). This is in line with previous observations that *C. albicans* telomeric and subtelomeric regions are highly dynamic (56–58). In contrast, centromeric regions did not display altered mutation rates relative to the genome average (Supplementary Figure 3C). This contrasts with the high mutation rates observed at *S. cerevisiae* centromeres (54), however the 3-4.5 kb regional centromeres present in *C. albicans* are much larger than the point centromeres (often <400 bp) found in this model yeast (59, 60).

Repeat sequences are common in fungal genomes and have been associated with mobile elements and gene regulation (61, 62). However, studies in model yeast have tended to exclude analysis of repeat regions to simplify genome analyses (40, 45). *C. albicans* is unusual among *Candida* species in that it contains 9 large Major Repeat Sequence (MRS) elements that span ~1.7% of the genome and are linked to chromosome translocations and chromosome length polymorphisms (62–66). We found that microevolved *C. albicans* lineages exhibited significant differences in mutation rates between repeat and non-repeat regions. For example, both MRS elements and long terminal repeat (LTR) retrotransposons showed mutation rates that were 13.1-fold higher than the genome average (Figure 3D and Supplementary Figure 3D). Importantly, a number of these mutations were validated by KASP analysis confirming that they arose during microevolution (Supplementary Table 4). Mutation rates in repeat regions were also significantly higher in strains passaged *in vivo* than *in vitro* (Supplementary Figure 3D).

Genes encoding GPI-linked cell wall proteins and *ALS* family genes are rich in internal tandem repeats that can vary in number and thereby contribute to allelic diversity and phenotypic variation (67–70). In line with these observations, we found that both GPI and *ALS* gene families accumulated mutations at much higher rates than the genome average (~8.6-fold increase, Figure 3D and Supplementary Figure 3E). These results establish that numerous chromosomal features including genomic repeats, telomeric regions, *ALS* genes and GPI-linked cell wall genes undergo evolution faster than the rest of the genome.

We also compared the distribution of mutations arising in heterozygous versus homozygous regions of the genome, while noting that LOH events in *C. albicans* can often promote adaptation (30, 71, 72). Heterozygous and homozygous regions were mapped in each of the four parental strains based on the density of heterozygous positions per 5 kb window (24). Using this metric, heterozygous regions in the four parental isolates varied between 69.5% and 84.2% of the genome (Supplementary Table 8). Comparison of the frequency of mutations between heterozygous and homozygous regions revealed that these regions accumulated mutations in line with their relative abundance (Figure 3D). For example, 84.2% of the P78048 genome is represented by heterozygous regions and 80.2% of all mutations in evolved derivatives of this lineage occurred in heterozygous regions. As LOH events are likely biased by the higher frequency of heterozygous positions in HET regions than in HOM regions, we repeated this analysis using only GOH events (base substitutions and indels). We again found that GOH events accumulated in HET regions at similar levels to their proportion in the genome (Supplementary Figure 3F). We therefore did not observe bias in the pattern of *de novo* mutations towards either heterozygous or homozygous regions of the genome.

### Microevolution is punctuated by frequent short-tract LOH events and small indels

A wide variety of LOH events have been described in *C. albicans*, with elevated LOH rates observed during exposure to stress, antifungal treatment, DNA damage, and host passage (22, 31, 32, 37, 73). To provide a global picture of these events during microevolution, we divided LOH events into three categories based on length (in kb) and the number of heterozygous positions affected: (1) microLOH (mLOH) events that involved loss of single heterozygous positions, (2) short-tract LOH (SLOH) events that involved loss of 2 or more heterozygous positions and covered small genomic regions (≤10 kb), and (3) long-tract LOH (LLOH) events that were >10 kb and affected hundreds of heterozygous positions (Figure 4A). The relative frequency of mLOH and SLOH events observed during microevolution was similar across experiments, with the minimum sizes of these events ranging between 1 and 3090 bp (L_min_ size, Figure 4B and Materials and Methods). Thus, isolates underwent an average of 52.3% mLOH and 46.4% SLOH events during passaging (Figure 4C,D). Analysis of the average size of LOH tracts revealed that L_avg_ varied between 222 bp (P57055) and 889 bp (SC5314) for the four strain backgrounds and impacted between 1.6 and 3.2 heterozygous positions (when excluding LLOH events, Supplementary Figure 4A,B). In contrast to frequent mLOH and SLOH events, long-tract LOH events occurred in only 2 passaged isolates (P76055 and P78048 grown in the GI SD model) and involved tracts of 273-1230 kb that extended to the ends of the chromosomes.

**Figure 4.**
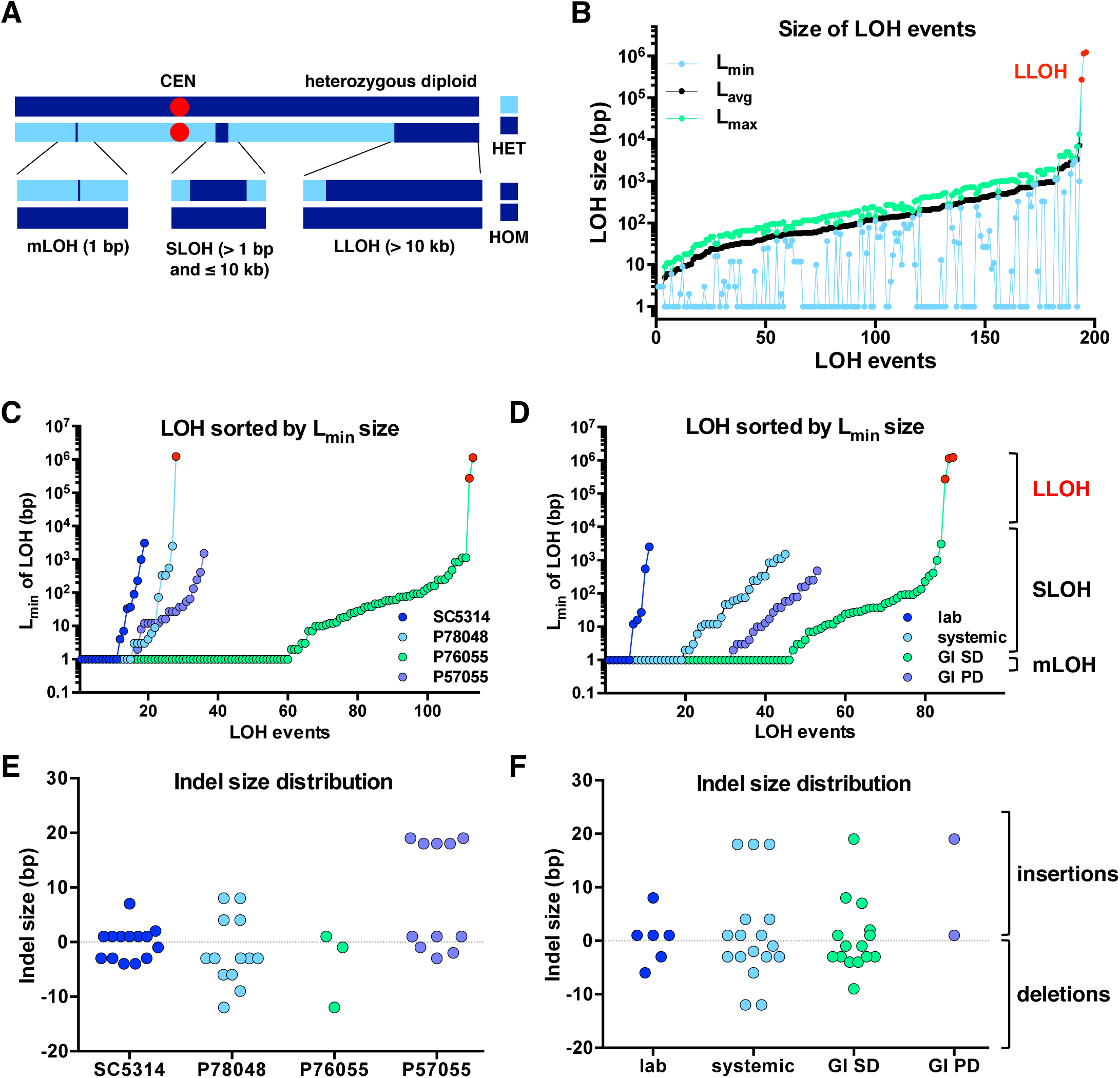
Microevolution is punctuated by frequent short-tract LOH events and small indels. (A) Schematic of different types of LOH events, including microLOH (mLOH, involve loss of single heterozygous positions), short-tract LOH (SLOH, involve loss of two or more heterozygous positions and are <10 kb), and long-tract LOH (LLOH, affect hundreds of heterozygous positions and are >10 kb). (B) Distribution of LOH events showing the L_min_, L_avg_ and L_max_ size for each LOH event. (C,D) L_min_ size distribution of LOH events, including microLOH, SLOH and LLOH (shown in red), for each lineage (C) and niche (D). (E,F) Size distribution of indels, including insertions and deletions for each lineage (E) and niche (F).

We also analyzed the 41 indels that occurred in the 28 evolved isolates (Figure 4E,F), finding that they were represented by similar numbers of insertions (21 events) and deletions (20 events). In *S. cerevisiae*, indels were biased towards insertions in haploid lines and towards deletions in diploid lines (40, 41). Indel sizes averaged 3.3 bp for *in vitro*-evolved isolates whereas indels were 2-3-fold larger for *in vivo*-passaged isolates (Figure 4F), although no significant differences in size were found between different lineages or different niches either for LOH events or for indels (*P* > 0.05). Together, these results provide the first comprehensive analysis of LOH and indel events in *C. albicans* and highlight that short-tract LOH events, most of which involve homozygosis of single heterozygous positions, are the most common LOH event occurring during microevolution.

### LOH events are overrepresented in repeat regions and telomeric regions

The frequency and distribution of LOH events arising within the *C. albicans* genome were examined for potential relationships with underlying chromosomal features. All LOH events (196 tracts) were mapped along the genome (Figure 5A) and their frequency determined per 0.2 Mb window (Supplementary Figure 4C). Mapping the distance of each LOH to the closest chromosomal feature revealed that a high proportion of LOH tracts (21%) arose either within MRS regions or in the 1 kb tracts adjacent to MRS or telomeric regions (Figure 5B and Supplementary Figure 5C). MRS elements were the main hotspot for these recombination events, with the start sites for 12% of all LOH tracts being located at these elements (Figure 5B). Not all MRS tracts showed the same propensity for recombination; MRS6 and MRS7b displayed the highest LOH frequencies and MRS2, 3, and 5 displayed the lowest LOH frequencies (Supplementary Figure 4C). In contrast, no LOH events were detected within 1 kb of the centromeres (Figure 5B). These findings establish that MRS and telomeric regions are hotspots for recombination in *C. albicans* and thereby promote genetic variation.

**Figure 5.**
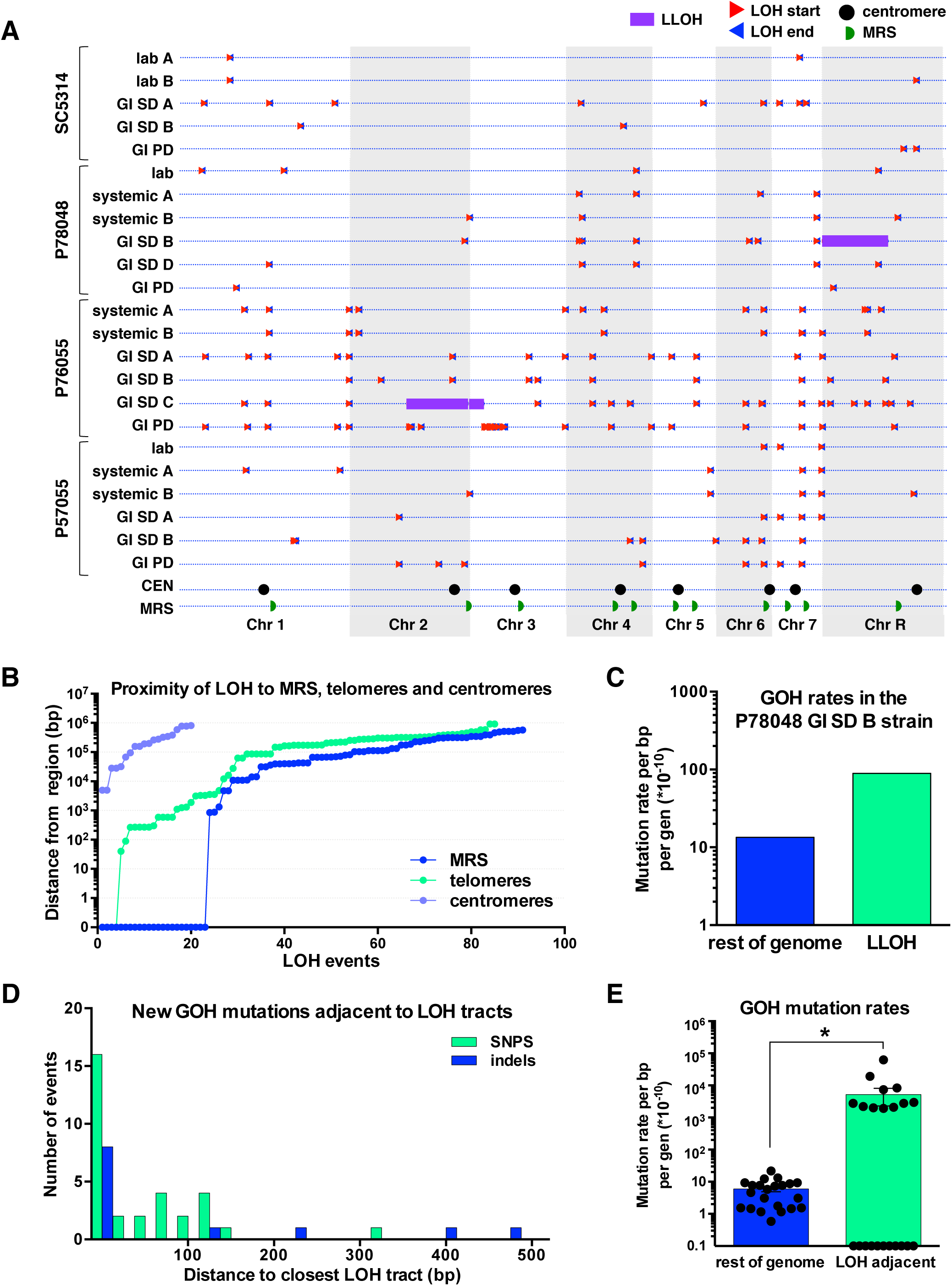
Relationship between LOH location and different genomic regions. (A) Chromosomal location of all LOH events (using L_min_) with triangles marking the start (red) and end (blue) of each event. Location of centromeres (CEN) and MRS regions are shown. (B) Proximity of LOH events to the closest genomic feature, including MRS regions, telomeres (Chr ENDs or *TLO* genes), and centromeres. Each LOH event is uniquely mapped to the closest of these features on the same chromosome arm. Distances equal to 0 indicate an LOH start site inside the respective genomic region. (C) GOH rates (including SNPs and indels) in the duplicated LLOH region versus the rest of the genome in the P78048 GI SD B isolate. Only GOH SNPs (base substitutions) were observed in the duplicated LLOH region. (D) Number of GOH events (SNPs and indels per 25 bp) observed within 500 bp of LOH tracts in microevolved isolates. (E) GOH rates (including SNPs and indels) in regions adjacent to LOH tracts (within 500 bp) compared to rates in the rest of the genome.

### The emergence of new heterozygous SNPs is detected within large LOH tracts

Three very large LOH tracts (0.27-1.23 Mbp) were formed during passaging, and these involved two isolates cultured in the GI (antibiotic-treated) model with the standard mouse diet (SD). LOH events involved the terminal regions of Chr 2 and Chr 3 in P76055 (GI SD C) and Chr R in P78048 (GI SD B; Figure 5A and Supplementary Figure 5A). LOH occurred via truncation of chromosome arms in P76055 GI SD C, as the resulting LOH tracts were monosomic (displayed half the read coverage for LOH regions on both Chr 2 and Chr 3, Supplementary Figure 1B). In contrast, LOH likely involved break-induced replication (BIR) or inter-homolog crossing-over in P78048 GI SD B, as the chromosome was still disomic for the emergent homozygous region (Supplementary Figure 1B). Analyses revealed that these large events led to LOH of thousands of heterozygous positions (as well as tens of indels) that were present in the parental genomes. In total, the three LLOH led to the homozygosis of 5,419 sites in coding regions, 2135 of which resulted in nonsynonymous changes, 14 produced nonsense mutations and three resulted in readthrough mutations (Supplementary Figure 5B and Supplementary Table 5). Interestingly, the LLOH region in P78048 GI SD B showed the reemergence of heterozygous positions due to *de novo* base substitutions (GOH SNPs) within the tract that had undergone LOH. In fact, a 6.6-fold higher rate of GOH events was detected within this LLOH tract than was evident in the rest of the P78048 GI SD B genome (Figure 5C). The high frequency of GOH mutations within the LOH tract is consistent with a high rate of *de novo* base substitutions emerging during BIR, as shown for *S. cerevisiae* where BIR was highly mutagenic (74).

### Impact of LOH events on mutational patterns

Chromosomal crossovers can induce *de novo* mutations in DNA regions close to the crossover in a wide variety of species (4, 50, 51, 75, 76). We therefore examined whether the regions flanking emergent LOH tracts in microevolved *C. albicans* isolates showed altered mutation rates relative to the genome average. Strikingly, sites adjacent to LOH tracts appeared highly enriched for mutations; 44 out of 136 GOH events (32 *de novo* base substitutions and 12 indels) were located within 500 bp of emergent LOH tracts (Figure 5D). In fact, 36 of these GOH events were located within just 100 bp of LOH tracts. Thus, 32.4% of all GOH events (26% of base substitutions and 92.3% of indels) were found in regions close to new LOH tracts, even though these regions represent only ~1.3% of the genome. GOH rates in LOH-adjacent regions (defined as 500 bp up/down of LOH tract) were therefore 840-fold higher than in the rest of the genome (Figure 5E), and were significantly higher in both systemic and GI SD infection models than in other environments (Supplementary Figure 5C). The distribution of GOH events adjacent to LOH tracts differed from that in the rest of the genome. For example, indels were highly enriched in LOH-adjacent regions, whereas GOH mutations arising in *TLO* genes, centromeric, and telomeric regions were not closely associated with LOH events (Supplementary Figure 5D).

We further note that *de novo* base substitutions in LOH-adjacent regions showed a *Ts/Tv* ratio of 1.13:1 compared to a genome average of 1.27:1. This is consistent with increased transversion rates resulting from translesion polymerases acting to repair DNA lesions at or close to recombination tracts (77, 78). In addition, both homologous recombination and non-homologous end-joining are considered to be error-prone mechanisms that can introduce indels close to the DNA break site (79). Our data now reveal that approximately a third of all GOH indels and substitutions in *C. albicans* arise in regions flanking LOH events and that this is likely due to highly mutagenic DNA repair mechanisms.

## Discussion

This study defines the spectrum of mutations that emerge in heterozygous diploid genomes of *C. albicans* during microevolution, including a comparison of mutational patterns during *in vitro* culture with those that occur during infection of a mammalian host. Numerous studies have established that *C. albicans* exhibits extensive genomic plasticity, from variation at the level of single-nucleotide polymorphisms to changes at the whole-chromosome level (24, 25, 30, 32, 64, 80). However, this work provides the first comprehensive picture of the genetic changes accompanying microevolution in this important pathogen.

### Global patterns of mutation in *C. albicans*

Microevolution resulted in similar mutational patterns regardless of the strain background or culture niche. In each case, microevolution was driven almost exclusively by multiple, small-scale changes in heterozygous polymorphisms (87.2%) and indels (12.8%). This reveals that ‘micro-scale changes’ are by far the most frequent events arising in the *C. albicans* genome. We establish that *C. albicans* displays an average *de novo* base-substitution rate of 1.17 × 10^−10^ per bp per generation during *in vitro* passaging. This is the first genome-wide estimate of *C. albicans* mutation rates and is close to those reported for mitotically dividing cells in the model yeast *S. cerevisiae* and *S. pombe* (8, 40, 41, 45). *C. albicans* therefore exhibits a *de novo* substitution rate similar to that of haploid or homozygous diploid yeast genomes. Critically, we show that microvariation in *C. albicans* is equally driven by LOH events, as these recombination events occur at frequencies (1.61 × 10^−10^ per bp per generation) that are close to those of *de novo* substitution rates and impact a similar number of nucleotide positions.

### Genome architecture and environmental pressures impact *C. albicans* microevolution

While overall mutational patterns were similar between microevolution lineages, *de novo* substitution and LOH rates varied significantly between different strain backgrounds and environments. For example, mutation rates varied by up to 5.6-fold between strains, although no obvious genetic differences were found (such as ‘mutator’ genotypes due to disruptions in DNA repair genes) that could account for these differences (see Supplementary Material). The niche in which strains were evolved had an even bigger impact on mutation rates; strains passaged *in vivo* showed up to 12.7-fold higher mutation rates than those passaged *in vitro*. *C. albicans* therefore experiences environment- or stress-induced mutagenesis as demonstrated for a number of bacterial, fungal, plant and human studies (81–83). In support of this, *C. albicans* was previously shown to undergo stress-induced LOH events *in vitro* (22), and certain long-tract LOH events were more frequent during bloodstream passage than during *in vitro* culture (31).

Our studies also establish that a number of chromosomal features impact *C. albicans* mutation rates. Mutation rates were higher in repeat regions, telomeric/subtelomeric regions, and in genes encoding GPI-linked cell wall proteins (including *ALS* family genes) than in the rest of the genome (see schematic in Figure 6). These results are consistent with multiple reports linking higher mutation rates within repetitive and telomere-proximal regions of the *C. albicans* genome (56, 62, 64, 67, 69, 84).

**Figure 6.**
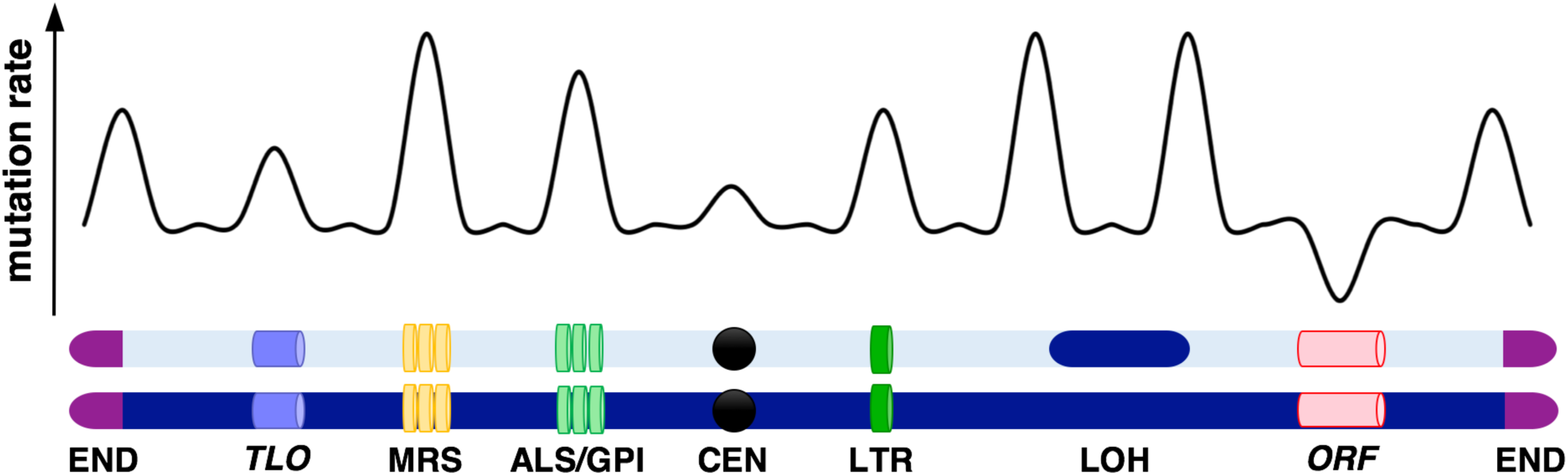
Schematic illustrating the pattern of mutational events across the *C. albicans* genome. Figure highlights how certain chromosomal features are associated with elevated mutation rates.

### Purifying selection acts on emerging mutations

Mutations accumulated at significantly higher rates in intergenic regions than in coding regions, and the ratio of synonymous to nonsynonymous mutations was also greater than that expected by chance. We estimate that 71-79% of nonsynonymous mutations were effectively removed from the population by selection based on the number of synonymous substitutions and the number of substitutions observed in intergenic regions. These results imply that purifying selection frequently acts to remove fitness-reducing mutants from the population. Natural isolates of *S. cerevisiae* also show a significant bias towards intergenic over genic SNPs (85), and more than a third of nonsynonymous mutations were implicated as being deleterious in one study (86). The current study provides striking evidence for purifying selection acting broadly on the diploid *C. albicans* genome even over relatively short evolutionary periods.

### Aneuploid forms frequently arise during passaging in one microevolution niche

Aneuploid forms have frequently been described in *C. albicans* (22, 24, 25, 87), yet only 3/28 microevolved lineages became aneuploid during our studies. In each case, diploid strains acquired a third copy of Chr 7 and these aneuploidies emerged in three different strain backgrounds during passaging in the GI tract using a standard mouse diet together with antibiotics. This suggests that being trisomic for Chr 7 provides a significant advantage under these growth conditions. To our knowledge, this is the first time this trisomy has been associated with passaging of *C. albicans* in the host. Interestingly, chromosome 7 trisomies did not emerge in isolates passaged in an alternative GI model that did not involve the use of antibiotics. This suggests that this trisomy does not enhance growth in the GI tract *per se* but could provide a specific advantage to attributes of the GI environment when mice are on the standard diet.

### MicroLOH events are a major driver of genome dynamics

LOH has long been recognized as an important mechanism for introducing diversity into *C. albicans* populations (21, 22, 24, 25, 30), although a global analysis of dynamic events had not previously been performed. Studies have examined the length of LOH tracts in diploid *S. cerevisiae* strains and showed that mitotic tracts are generally longer than meiotic tracts, with the former averaging 2 - 12 kb (88, 89). A recent study examined genome-wide recombination events in mitotic *S. cerevisiae* cells and showed that LOH tracts range from <100 bp to >100 kb with small LOH tracts (<1 kb) attributed to local gene conversions, although many of the smallest LOH events were excluded from this analysis (8). The current study now defines the total spectrum of LOH events occurring during *C. albicans* microevolution. *C. albicans* LOH rates were 1.61 × 10^−10^ per bp per generation (or 4.5 × 10^−3^ per cell division) which are slightly lower than previous estimates of 1.3 - 2.9 × 10^−9^ per bp per generation based on events at three select loci (66, 90). However, we note that LOH rates varied considerably between different genomic regions, with MRS and telomeric regions representing relative hotspots for LOH.

We reveal that the majority of LOH events in *C. albicans* involve very short microLOH tracts (mLOH, estimated L_avg_ size = 368 bp). Indeed, over half (52%) of all LOH events impacted only a single heterozygous position and, critically, a number of these events were validated by KASP genotyping. In fact, when examining all genetic changes accrued during microevolution, >30% of these changes were due to LOH at single heterozygous positions revealing that these represent a very high frequency event in the *C. albicans* genome. Consistent with our data, experiments studying the repair of DNA double-strand breaks in *C. albicans* showed frequent short-tract LOH events via gene conversion, and only rarely were long-tract LOH events observed due to BIR (Break-Induced Replication) or reciprocal recombination (73). Similarly, recent analysis of passaged lineages in the water flea *Daphnia pulex* also found that short-tract LOH events (median ~221 bp) were prevalent (7). We therefore suggest that short-tract LOH events represent a common occurrence in heterozygous diploid genomes but will have been missed by studies that lack nucleotide-level resolution.

In contrast to frequent microLOH events, large LOH tracts were rarely observed in our experiments and involved only two isolates passaged in the GI SD model. These LOH events involved long DNA tracts (0.27-1.23 Mb) that extended to the ends of the chromosomes. Previous studies also detected large LOH events in *C. albicans* strains grown both *in vitro* and *in vivo* (21, 22, 24, 25, 73, 91). We note that while large-scale chromosomal changes are relatively rare, these impact a large number of genes and are therefore the most likely to have phenotypic consequences. Overall, the three large LOH events identified here led to homozygosis of over 10 thousand heterozygous positions, resulting in 2,135 nonsynonymous changes, 14 nonsense mutations and 3 readthrough mutations. This reveals that large LOH tracts drive extensive genotypic changes but may also be heavily selected against given the large number of positions impacted by such events.

We were also surprised to find that global LOH rates were balanced by equivalent rates of *de novo* GOH mutations during microevolution, regardless of genetic background or evolution niche. Because of this balance, overall heterozygosity levels were stably maintained (± 2%) in the majority of passaging experiments. This finding sheds light on an important question in *C. albicans* – how do strains maintain genome heterozygosity levels in the face of frequent LOH events? The observation that *de novo* mutation rates often match LOH rates indicates that genomes can frequently remain heterozygous even in the absence of outcrossing events.

### An association between recombination events and *de novo* mutations

We found that there was a striking correlation between *de novo* mutations (both base substitutions and indels) and their proximity to recombination events in *C. albicans*. Specifically, GOH rates were elevated 840-fold within 500 bp of emergent LOH tracts relative to the genome average. Consequently, a third of all GOH events (substitutions and indels) were in regions flanking new LOH tracts, despite these tracts representing only ~1.3% of the genome. This phenomenon was observed independent of strain background and was most evident during host passage where LOH rates were higher. The high rate of mutations adjacent to LOH tracts is likely due to LOH being mutagenic and introducing *de novo* mutations into neighboring regions of the genome. This is consistent with DNA double-strand break repair processes being both recombinogenic and mutagenic, as has been described in several cell types (51, 74, 92). Increased rates of transversions in these mutated regions support the activity of translesion polymerases acting to repair DNA lesions at these sites. However, this is the first evidence that recombination events in *C. albicans* introduce *de novo* mutations during the repair process. Thus, LOH is a stress-inducible event in *C*. *albicans* (*22*) and also introduces additional mutations into the genome, both of which will accelerate adaptation.

### Concluding remarks

This study provides a high-resolution analysis of the spectrum of mutations accumulating in a heterozygous diploid pathogen. We demonstrate that both cell-intrinsic properties (e.g., strain background, repetitive chromosomal features) and cell-extrinsic factors (e.g., *in vivo* versus *in vitro* passage) impact the frequency and distribution of genetic fluctuations. Frequent micro-scale changes (predominantly *de novo* substitutions and short-tract LOH events) and occasional larger-scale rearrangements (long-tract LOH or chromosomal aneuploidies) determine genome dynamics. Furthermore, purifying selection plays a dominant role in dictating which genetic changes are retained during evolution. Our results provide a detailed picture against which genomic changes in other heterozygous diploid species can be evaluated, and establish the foundation for understanding how *C. albicans* can adapt to a wide variety of distinct host niches.

## Acknowledgements

We would like to thank Adrian Vladu for help with computational analyses, and Daniel Weinreich and members of the Bennett lab for feedback on the manuscript. This work was supported by National Institutes of Health grants AI081704, AI122011 and AI112363 (to R.J.B.), by a PATH award from the Burroughs Wellcome Fund (to R.J.B.), by a Vessa Notchev Fellowship from Sigma Delta Epsilon-Graduate Women in Science (to I.V.E.), by a Wellcome Trust / MIT Postdoctoral Fellowship (to R.A.F), by a NIH F31 Pre-doctoral Fellowship (F31DE023726 to M.P.H.) and with Federal funds from the National Institute of Allergy and Infectious Diseases, National Institutes of Health, Department of Health and Human Services, under Contract No.:HHSN272200900018C and Grant Number U19AI110818 to the Broad Institute.

## Author contributions

I.V.E. and R.J.B. planned the experiments, I.V.E. and M.P.H performed the experiments, I.V.E, R.A.F., K.A. and C.A.C. performed the bioinformatics analyses. I.V.E and R.J.B. drafted the manuscript with contributions from R.A.F. and C.A.C.

## Materials and Methods

### Strains and growth conditions

The *C. albicans* strains used in this study are listed in Supplementary Table 1. Unless otherwise stated, strains were grown at 30°C in YPD medium (93). For *in vitro* evolution experiments, cultures were serially diluted (1/100) every day for 80 days (bottlenecks every 6.8 generations) and cells collected once a week. *In vitro* isolates were collected as a pool and used to prepare genomic DNA.

### Determination of doubling time and generation times

For determination of doubling times *C. albicans* strains were grown in YPD at 30°C for 18-24 h and cell densities were recorded every 10-15 min in a Biotek Synergy HT plate reader/incubator. Exponential growth intervals were selected for doubling time estimates. Doubling times (D) were calculated using the formula D = t / log_2_(N_t_/N_0_) with t = duration of growth interval, N_0_ = number of cells at the start of the selected interval and N_t_ = number of cells at the end of the selected interval. The four original starting strains displayed similar doubling times (~1.5 h, Supplementary Table 3). The number of generations during *in vitro* microevolution experiments was calculated using the starting and final cell densities as G = log_2_(N_t_/N_0_) with N_0_ = number of cells at the start of the culture and N_t_ = number of cells at the end of the culture (~6.8 generations/day, Supplementary Table 3). The reported doubling and generation times represent the averages of three biological replicates (with 2 technical replicates performed for each biological replicate).

### Murine experiments

For animal infections, 6-7-week-old female BALB/c mice (~18 g) from Charles River Laboratories were housed together with free access to food and water. For systemic infection, *C. albicans* cells were grown overnight in YPD medium at 30°C, washed in phosphate-buffered saline (PBS), and for each passage 3 mice were infected via the tail-vein with a total inoculum of 6 × 10^5^ cells in a 200 µl volume. Mice were monitored for signs of infection and their weights, posture and motility scored daily. 3 days post-infection the mice were euthanized and fungal cells isolated from infected kidneys using a PBS solution supplemented with an antibiotic mixture (500 µg/mL penicillin, 500 µg/mL ampicillin, 250 µg/mL streptomycin, 225 µg/mL kanamycin, 125 µg/mL chloramphenicol, and 125 µg/mL doxycycline). The number of colony forming units (CFUs) from the kidneys was determined by plating cells onto YPD medium. Isolates for subsequent passages were selected by picking single colonies from the mouse showing the highest virulence outcome score. Virulence outcome scores were determined by assessing kidney fungal burdens and weight changes at 72 h using the formula: outcome score = log (kidney CFU/g) − (0.5 × percentage weight change) (94). After five 3-day passages, the last isolates were collected, mice were humanely sacrificed and fungal burdens determined from the kidney. Colonies from the mouse showing the highest outcome score at passage 5 were used to prepare genomic DNA.

For GI colonization experiments, two different murine models of commensalism were used. For the standard diet (SD) model, mice were fed standard rodent chow (FormuLab 5001, PMI Nutrition International) and their water was supplemented with antibiotics (1500 units/mL of penicillin, 2 mg/mL of streptomycin) and 5% glucose for taste (95). The antibiotic treatment was initiated 4 days prior to infection to reduce the endogenous gastrointestinal microbiota. Alternatively, mice were fed a purified diet (PD) starting 4 days prior to inoculation in which case their water was not supplemented with antibiotics (34). In both models, mice were orally gavaged with a 20G × 38 mm plastic feeding tube (Instech Laboratories, Inc.) with 10^8^ *C. albicans* cells in a 500 µl volume and continued with their respective diet and water for 6 weeks. To prevent contamination between independent evolution experiments, each mouse was housed in a separate cage. Fecal samples were collected weekly and fungal cells isolated using a PBS solution supplemented with antibiotics. At 42 days, isolates were collected as a pool and used to prepare genomic DNA. After the last isolate collection, mice were humanely sacrificed and fungal burdens determined from the GI organs (stomach, small intestine, colon and caecum).

### Whole-genome sequencing and variant identification

To extract genomic DNA, *C. albicans* isolates (Supplementary Table 1) were grown overnight in YPD at 30°C and DNA isolated from ~10^9^ cells using a Qiagen Genomic Buffer Set, a Qiagen Genomic-tip 100/G or the MasterPure Yeast DNA Purification kit (Epicentre). Each isolate was sequenced using Illumina HiSeq 2000 generating 101 bp paired reads. The nuclear genome sequences and General Feature Files (GFF) for *C. albicans* SC5314 reference genome (version A21-s02-m08-r01) were downloaded from http://www.candidagenome.org/. We randomly down-sampled the paired-end Illumina reads for isolate SC5314 from 13 SRA runs (SRR1106648; SRR1106646; SRR1106647; SRR1106651; SRR1106653; SRR1106654; SRR1106656; SRR1106658; SRR1106664; SRR1106643; SRR1106645; SRR1106649; SRR1106655) to 45,130,695 paired reads (~300X deep, where the range for all isolates is 70X - 547X). Reads were aligned to the SC5314 reference genome assembly using Burrows-Wheeler Aligner (BWA) v0.7.4-r385 mem (96), and converted to sorted BAM format using Samtools v0.1.9 (r783) (97). The Genome Analysis Toolkit (GATK) (98) v2.7-4-g6f46d11 was used to call both variant and reference bases from the alignments. Briefly, the Picard tools (http://picard.sourceforge.net/) AddOrReplaceReadGroups, MarkDuplicates, CreateSequenceDictionary and ReorderSam were used to preprocess the alignments. We used GATK RealignerTargetCreator and IndelRealigner for resolving misaligned reads close to indels on parental-progeny pairs of isolates to avoid discrepancies between isolates. Next, GATK Haplotype Caller and Pilon (36) (with diploid genotyper ploidy setting) were run with both SNP and INDEL genotype likelihood models (GLM). We then merged and sorted all the calls from Haplotype Caller, and ran VariantFiltration with the following filters “QD < 2.0, FS > 60.0, MQ < 40.0, MQRankSum < −12.5, ReadPosRankSum < −8”. Next, we removed any base that had less than a minimum genotype quality of 50, or a minimum depth of 20. Finally, we removed any positions that were called by both GLMs (i.e., incompatible indels and SNPs), any marked as “LowQual” by GATK, nested indels, or sites that did not include a PASS flag. Similar filtering was performed for Pilon calls, removing low quality sites and setting a minimum depth of 20. All mutations in evolved isolates were visually inspected using IGV (http://software.broadinstitute.org/software/igv/). Identical mutations that were present in multiple isolates from the same lineage were removed from the analyses under the likelihood that they had been present in the parental strains. The final base calls covered >97% of the genome for any given isolate (Supplementary Table 2). We then categorized every single base between a parent and progeny (summarized in Supplementary Table 2), and annotated those changes using the GFF (VCFannotator, Broad Institute).

### KASP genotyping

To validate sequence variants, genomic DNA was subjected to allele specific PCR (KASP genotyping technology, LGC group), a fluorescent technique which enables testing of SNPs and indels at specific loci. Primers were designed with 50 bp flanks around the site of interest for each variant allele and genomic DNA from original and microevolved isolates was tested across 80 unique sites (20 for each strain background). Allele frequencies were calculated for each site and genotyping was assigned by cluster analysis. Sites were selected so that they represent all mutation categories (GOH, LOH, SNPs, indels, transitions, transversions) as well as different regions of the genome (coding, intergenic, MRS, retrotransposons, adhesins and telomeric genes). Out of the 80 SNPs tested, 63 (78.8%) were successfully genotyped via KASP. The 63 genotyped sites were then compared with mutations called from genome sequencing data with an 87.3% success rate.

### Ploidy and copy number variation

To examine ploidy variation across the genome, the Illumina read alignment depth was calculated for 100 bp windows across the genome, using BEDTools 2.18 (99), SAMtools 1.3 (97) and the GATK 3.7 Depth of Coverage module. The read depth was calculated as the number of bases aligned per window divided by the length of the window and normalized to the average depth for each strain and to the GC content, as this can influence both the sequencing chemistry and the alignment quality (100). The read depth was also normalized per the effective window length by removing any ambiguous sites in the respective window. The normalized alignment depth for each 100 bp window was then plotted and large scale variations in ploidy (2 fold up or down coverage) were identified. These include whole chromosome and segmental aneuploidies larger than 0.1 Mbp. Smaller regions showing read depth variation were designated as copy number variants (CNVs) and their numbers plotted based on the nature of the variation (2 fold up or down coverage).

### LOH analysis

To identify LOH events, all variants were classified based on how each mutation alters heterozygosity at the respective site: losses of heterozygosity (LOH), gains of heterozygosity (GOH), or mutations that do not alter heterozygosity (het neutral). LOH tracts were defined using each heterozygous site identified to have undergone LOH and the size of the tracts was determined by visual inspection in IGV. To calculate L_min_ for an LOH event, tracts began at the first converted LOH SNP identified and ended at the final converted LOH SNP with no interruption by a heterozygous position. For LOH events encompassing a single LOH SNP (i.e., not flanked by a consecutive LOH SNP) L_min_ was 1. To calculate L_max_ for an LOH event, tracts measure the distance between the nearest upstream, non-converted position and the nearest first downstream, non-converted position of the respective LOH tract. The average size of LOH tracts (L_avg_) was calculated by averaging the minimum (L_min_) and maximum (L_max_) lengths for each observed event. LOH tracts were then classified based on genomic size: microLOH (mLOH, affecting single heterozygous positions and with an L_min_ of 1 bp), short tract LOH (SLOH, affecting two or more heterozygous positions and with an L_min_ ≤ 10 kb), and long tract LOH (LLOH, affecting hundreds of heterozygous positions and with an L_min_ > 10kb). The numbers of LOH were then assessed for each lineage, niche of evolution and chromosome. The LOH distribution was examined across all isolates for each lineage and for each niche using the L_min_ genomic size (Figure 4C,D) of individual LOH events or the number of heterozygous positions that were impacted by each LOH (Supplementary Figure 4B).

### Mutations in different genomic regions or regions adjacent to LOH events

We identified mutations in specific regions using the genomic coordinates of these regions - strain specific HET (heterozygous) and HOM (homozygous) regions, HET/HOM junctions, repeat regions (MRS, LTR and genes associated with repeats), centromeres, chromosomal ends, *TLO* genes, and *ALS* and GPI-linked genes. Genomic coordinates for these chromosomal features were obtained from the Candida Genome Database (http://www.candidagenome.org) and genomic coordinates are provided in Supplementary Tables 6 and 7. HET and HOM regions were previously defined for the four starting strains (24) and are included in Supplementary Table 8.

### Statistical analyses

Statistical analyses were performed using two-tailed Student’s t-tests and by calculating probability values of binomial model distributions using Microsoft Excel 2016 (Microsoft) and Prism 6 (GraphPad). Significance was assigned for *P* values < 0.05, and asterisks denote *P* values that satisfy this condition.

### Data access

The sequence data from this study have been submitted to the NCBI SRA under BioProject ID PRJNA345600 (http://www.ncbi.nlm.nih.gov/bioproject).

### Ethics Statement

This study was carried out in strict accordance with the recommendations in the Guide for the Care and Use of Laboratory Animals as defined by the National Institutes of Health (PHS Assurance #A3284-01). Animal protocols were reviewed and approved by the Institutional Animal Care and Use Committee (IACUC) of Brown University. All animals were housed in a centralized and AAALAC-accredited research animal facility that is fully staffed with trained husbandry, technical and veterinary personnel.

## Supplemental Material

**Supplementary Text 1. Analysis of copy number variation during microevolution**

Read depth analysis revealed copy number variation (CNV) during microevolution. Several genomic regions (average of ~2% of all 100 bp windows) showed a two-fold increase or decrease in coverage relative to the parental strain. Both the number of CNV regions and the nature of the variation differed between strain backgrounds (Supplementary Figure 6A). For example, the SC5314 lineage showed the least CNV, with only ~0.2% of 100 bp windows displaying a two-fold decrease (2X down) relative to the starting isolate. In contrast, ~0.7% of the P57055 windows displayed a two-fold decrease in coverage and ~1.7% of P57055 windows displayed a two-fold increase (2X up) in coverage (Supplementary Figure 6A).

A significant proportion of CNV was associated with specific regions of the genome. Analysis focused on the major repeat sequences (9 MRS elements span 1.7% of the genome and have been linked to chromosome translocations (62–66)), as well as the terminal 5 kb regions of chromosomes and the *TLO* family of subtelomeric genes (56, 62, 64, 66, 101) (see Supplementary Tables 6 and 7 for genomic coordinates). In each of these regions, evolved isolates displayed a higher percentage of windows with variable coverage relative to the rest of the genome (Supplementary Figure 6B-D). Chromosomal ends and *TLO* genes more often displayed a higher number of windows with increased read coverage following passaging (Supplementary Figure 6B,C, and Supplementary Figure 7A,B), whereas MRS regions often displayed reduced coverage (Supplementary Figure 6D). This suggests that terminal regions and *TLO* genes often underwent expansion during microevolution experiments whereas MRS regions more often underwent contraction. A closer inspection revealed that decreased coverage (2X down) was equally observed across MRS subunits, whereas increased coverage (2X up) was most common in the RB2 subunit (Supplementary Figure 6E,F). This result was surprising as variation was expected to be greatest in the tandem array of highly repetitive RPS subunits (62). While the RPS regions displayed the highest coverage across the MRS (suggestive of multiple repeats being present), they displayed little CNV during our microevolution experiments (Supplementary Figure 7C).

We also note that CNV occurred between genes from different lineages, which could contribute to phenotypic differences between strains. For example, clade I strains displayed higher relative coverage of orf19.5474, encoding a protein of unknown function that is induced by Mnl1 during acid stress (102) (Supplementary Figure 7A). In contrast, this gene displayed lower coverage levels in the other 2 lineages. Similarly, variation in the relative coverage levels of *TLO* genes was noted between lineages (Supplementary Figure 7B). These likely represent variations in gene copy number but local effects of accessibility to DNA isolation and purification could also contribute to these differences. Overall, these analyses indicate that extensive copy number differences exist between different isolates of *C. albicans*. In addition, particular regions of the genome are more prone to CNV, including both the terminal chromosome and MRS regions which fluctuate between 0 and 40 copies, and the *TLO* genes which fluctuate between 0 and 8 copies (Supplementary Figure 7).

**Supplementary Text 2. Impact of mutations in DNA repair and maintenance genes on microevolution.**

To examine whether preexisting mutations could affect differences in mutation rates between lineages, we screened for mutations in ~60 genes involved in DNA maintenance and repair (Supplementary Table 9). This screen identified nonsense and readthrough mutations in several of the starting strains (Supplementary Table 10). For example, both clade I strains SC5314 and P78048 had 2 premature heterozygous stop codon mutations in *ARP8*, a predicted component of the chromatin-remodeling enzyme complex (103). These strains also had a heterozygous readthrough mutation in *FPG1*, a DNA glycosylase involved in the repair of irradiated DNA (104), whereas P76055 had a homozygous readthrough mutation in *FPG1* (Supplementary Table 10). The position of the *FPG1* mutations was identical between strains, indicating that they shared a common ancestor and that the readthrough mutation occurred before clade I and II diverged. In addition, P78048 had a unique heterozygous nonsense mutation in *SPO11*, a DNA endonuclease implicated in genetic recombination during parasex (105). As most of these mutations were heterozygous or present in genes that are not essential for DNA maintenance under our experimental conditions, it is likely that they did not play a major role in impacting mutation rates during microevolution.

We also examined whether new mutations emerged which could alter the function of DNA repair genes during passaging. Overall, the 28 isolates acquired a total of 128 mutations in genes with known or predicted roles in DNA maintenance or DNA repair, 120 of which were directly associated with large chromosomal events such as LLOH tracts (Supplementary Table 11). The 120 mutations associated with LLOH tracts involved loss of heterozygosity mutations indicating that they were a direct consequence of the LOH event (Supplementary Table 11).

The remaining mutations included 2 nonsynonymous changes (LOH mutations) in the *RAD50* gene of the P76055 GI PD isolate, which encodes a double strand repair protein with roles in stress responses (106). However, none of the identified mutations were associated with obviously increased mutation rates in the respective isolates (Figure 3A). Mutations in *RAD57* (orf19.4275) in one isolate (P76055 GI SD C) included 4 nonsynonymous missense mutations (Supplementary Table 11). These mutations were in the first third of the amino acid sequence and could disrupt function. *RAD57* encodes a key protein involved in DNA recombination and these mutations possibly altered the ability of this isolate to successfully undergo BIR and return to normal diploid levels, as illustrated by copy number variation analyses (Supplementary Figure 1B). Therefore, large-scale chromosomal changes via LOH events may have long-term consequences for genome evolution by disrupting important DNA repair pathways.

## Supplementary Figure legends

**Supplementary Figure 1.**
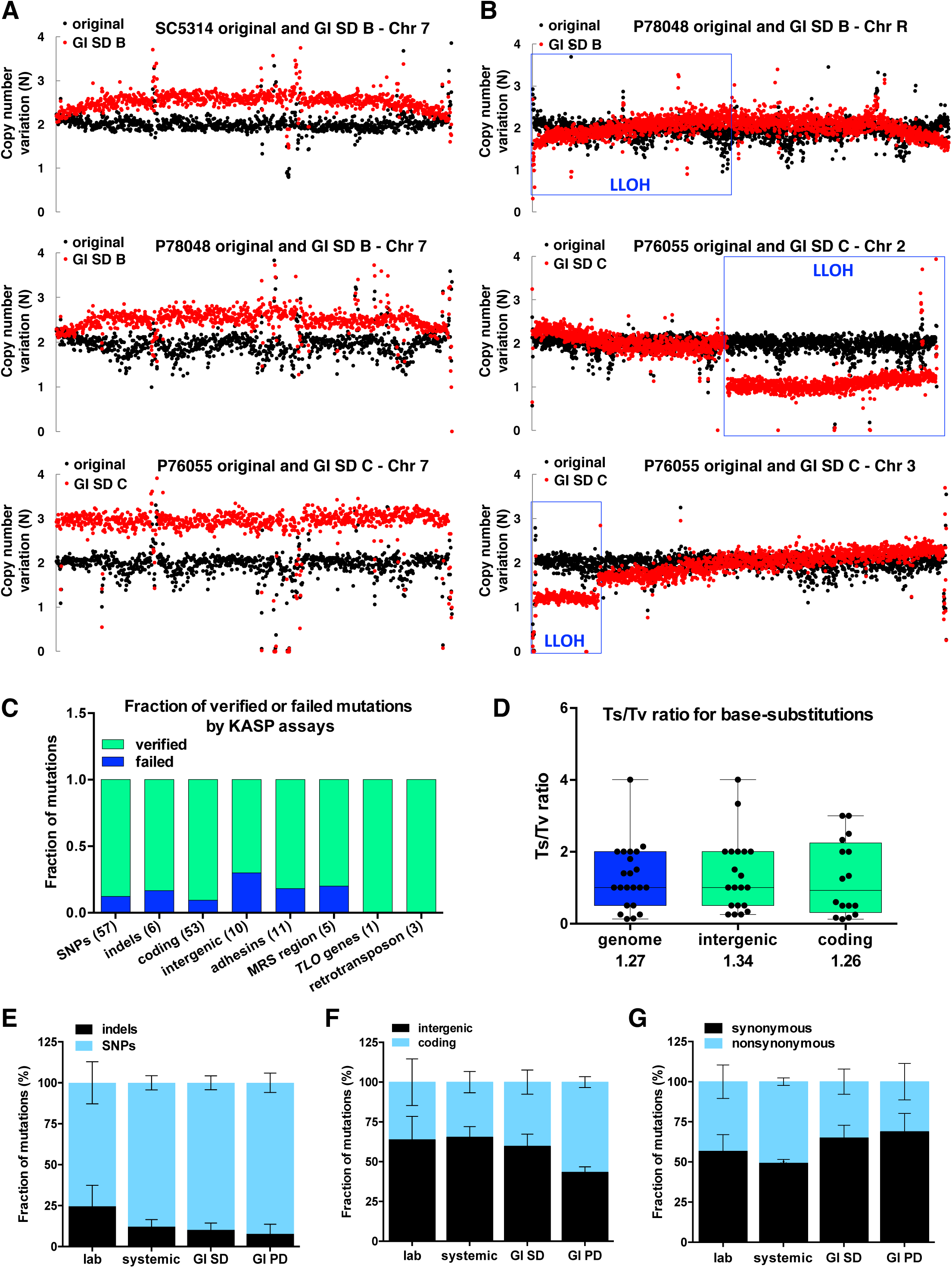
Large chromosomal events identified during microevolution. (A) Chr 7 trisomies were present in isolates recovered from the GI (SD model) in lineages SC5314, P78048 and P76055. (B) Segmental aneuploidies for Chr 2 and 3 were present in isolate P76055 GI SD C (boxed). A third LLOH tract was identified on Chr R of P78048 GI SD B (boxed). Variations in ploidy were determined by calculating the normalized read depth per 1 kb window. (C) Mutations verified using KASP assays. Fractions show the number of assays that failed or verified events identified via Illumina sequencing for different types of mutations. (D) Transitions/Transversion (*Ts/Tv*) ratios calculated for mutations arising during microevolution and broken down for mutations identified in intergenic and coding regions. Average *Ts/Tv* ratios are included below each category. (E) Distribution of SNPs and indels, coding and intergenic mutations, and synonymous and nonsynonymous mutations across microevolved isolates averaged for each niche. Note that panels include both GOH and LOH events.

**Supplementary Figure 2.**
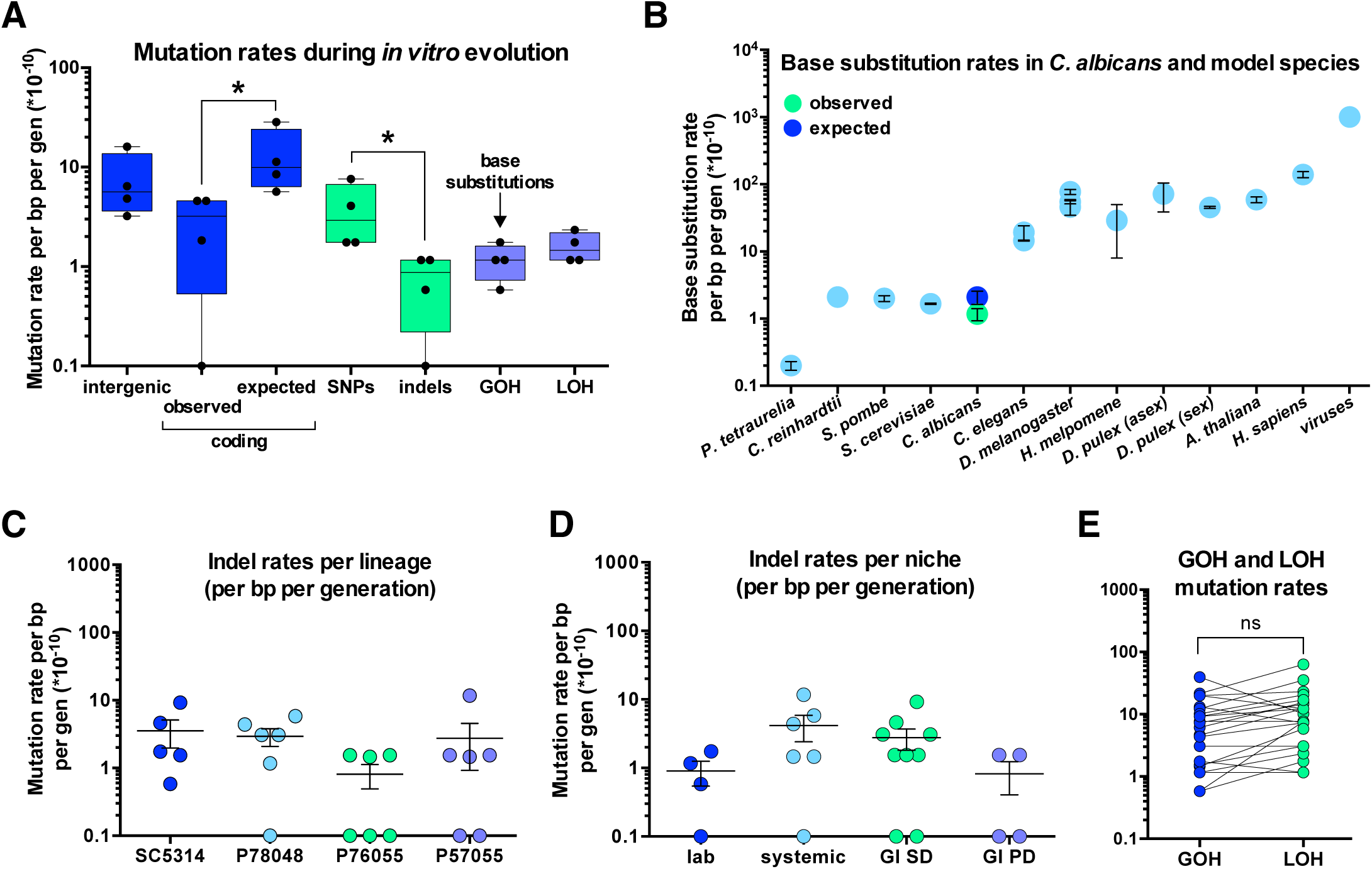
(A) Mutation rates (per bp per generation) calculated for different types of mutations following *in vitro* passaging of isolates. Asterisks indicate significant differences (t-test, *P* < 0.05). (B) Comparison of base-substitution rates in *C. albicans* and model organisms. Base-substitution rates (GOH SNPs) for *C. albicans* are shown as an average from *in vitro* evolution experiments performed in four different experiments and three genetic backgrounds. Expected rates reflect estimates based on observed intergenic base-substitution mutations. (C,D) Effect of strain background (C) and evolution niche (D) on indel mutation rates. Indels shown are the result of both GOH and LOH events. (E) GOH and LOH mutation rates across microevolution experiments. No significant difference was found between the two groups (t-test, *P* < 0.05).

**Supplementary Figure 3.**
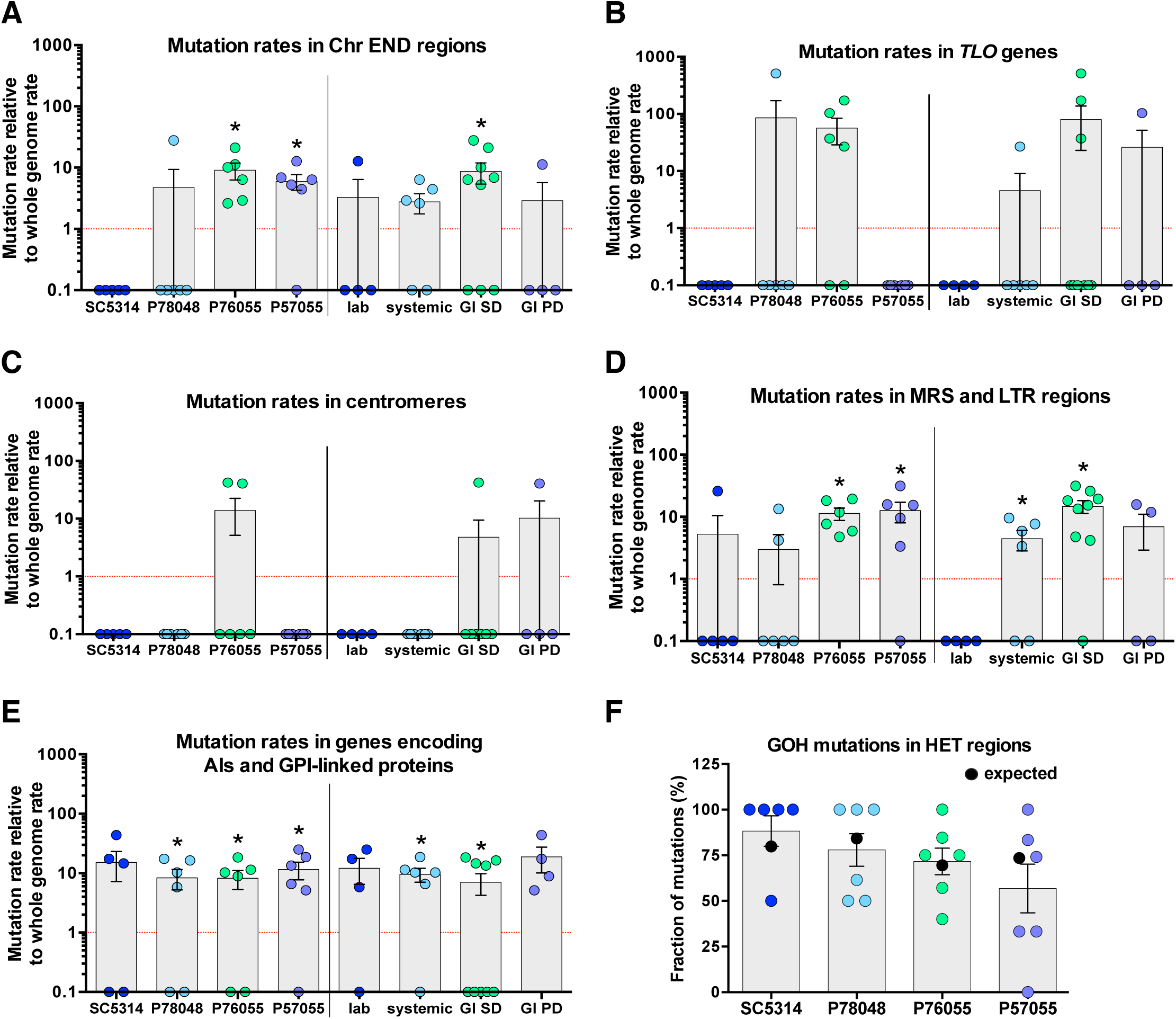
Mutation rates in specific regions of the *C. albicans* genome. Panels show mutation (SNPs and indels due to both GOH and LOH) rates relative to whole genome rates for (A) Chr END regions (final 10 kb of each Chr arm), (B) *TLO* genes, (C) centromeres, (D) repeat regions (MRS and LTR), and (E) genes encoding *ALS* family proteins and GPI-linked proteins. Asterisks indicate significant differences relative to either the SC5314 lineage or *in vitro* passaged isolates (*P* < 0.05). (F) Observed fraction of GOH mutations in heterozygous (HET) regions for each lineage. Expected values (black) represent the % HET regions in the parental isolates of each lineage based on the density of heterozygous sites per 5 kb genomic windows.

**Supplementary Figure 4.**
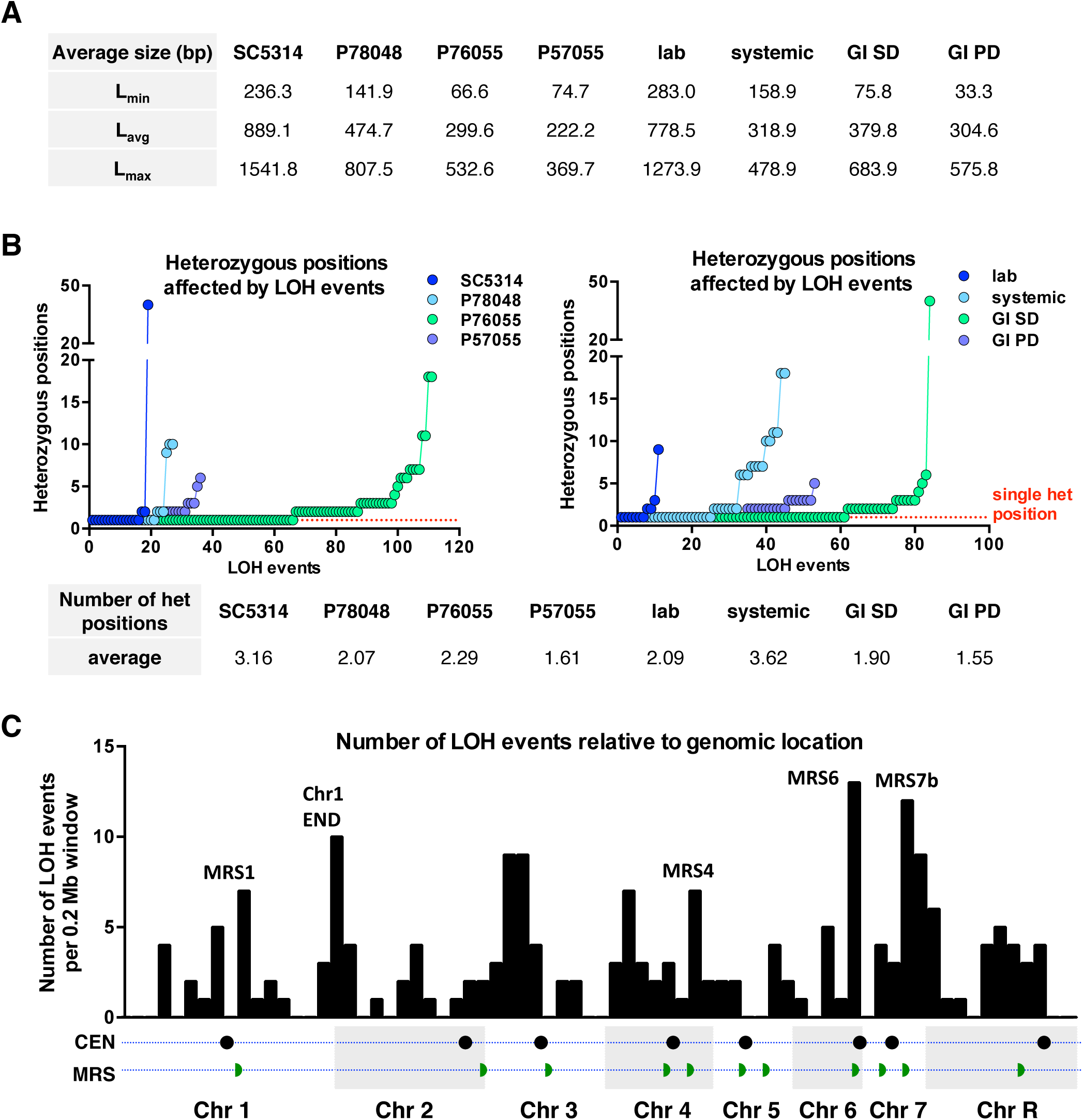
Size of LOH events arising during microevolution. (A) Average L_min_, L_avg_ and L_max_ size of LOH events for each lineage and niche. (B) Number of heterozygous positions affected by LOH tracts, shown for each lineage and niche. The average number of heterozygous positions impacted by LOH are included for each lineage and niche. For panels A and B, the three large LOH events were excluded from the analyses. (C) Density of LOH events relative to genomic location (per 0.2 Mb windows). Centromeres and MRS regions are included for reference.

**Supplementary Figure 5.**
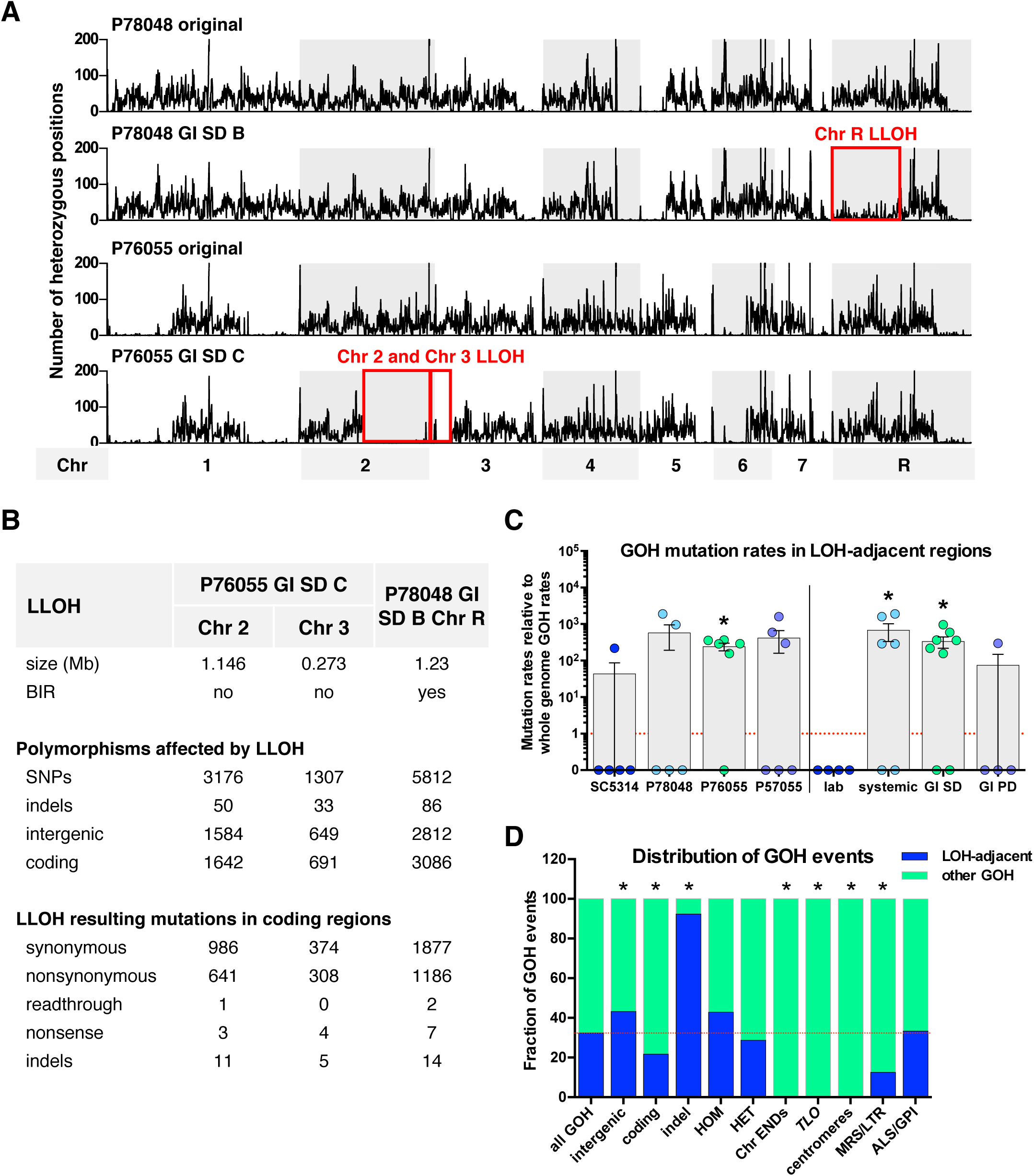
LLOH events represent three contiguous large-tract LOH regions. (A) Heterozygosity plots indicating the number of heterozygous positions for each 10 kb window across the 8 *C. albicans* chromosomes. Large LOH tracts on Chr R, 2 and 3 are boxed in red and shown relative to corresponding parental isolates. (B) Genomic size of tracts and number of mutations resulting from LLOH events, including a breakdown for mutations in the coding region. (C) GOH mutation rates in the 500 bp regions flanking LOH events (upstream and downstream) relative to whole genome GOH rates. Asterisks indicate significant differences (t-test, *P* < 0.05) relative to whole genome rates (red dotted line). (D) Distribution of GOH events (LOH-adjacent or not) relative to their position in the genome. Regions showing enrichment of one GOH category over another are marked with asterisks (*P* < 0.05, using a binomial distribution model).

**Supplementary Figure 6.**
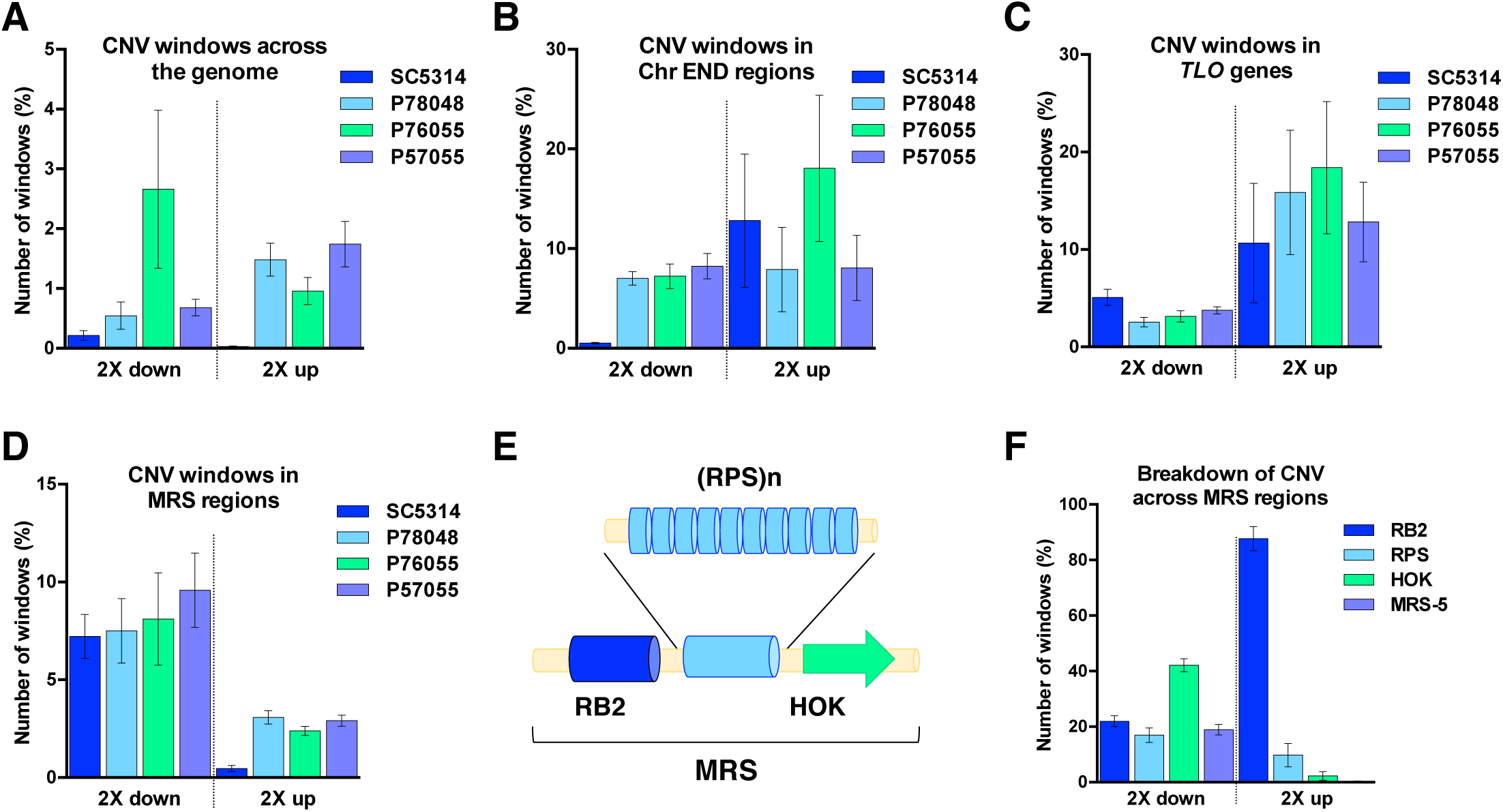
Copy number variation (CNV) analysis. (A) Summary of CNV windows across the genome showing a two-fold increase (2X up) or decrease (2X down) in coverage relative to the original strain and averaged for each of the four lineages. (B-D) Summary of CNV windows showing a two-fold increase or decrease relative to the original strain in Chr END regions (terminal 5 kb, B), *TLO* genes (C) and MRS regions (D). (E) Schematic representation of a typical MRS region, including RB2, RPS and HOK subregions. (F) Breakdown of CNV windows aligning to different MRS subregions or MRS-5 (only partial MRS sequences are present on Chr 5) (63). For all analyses CNV was determined by calculating the normalized read depth per 100 bp window.

**Supplementary Figure 7.**
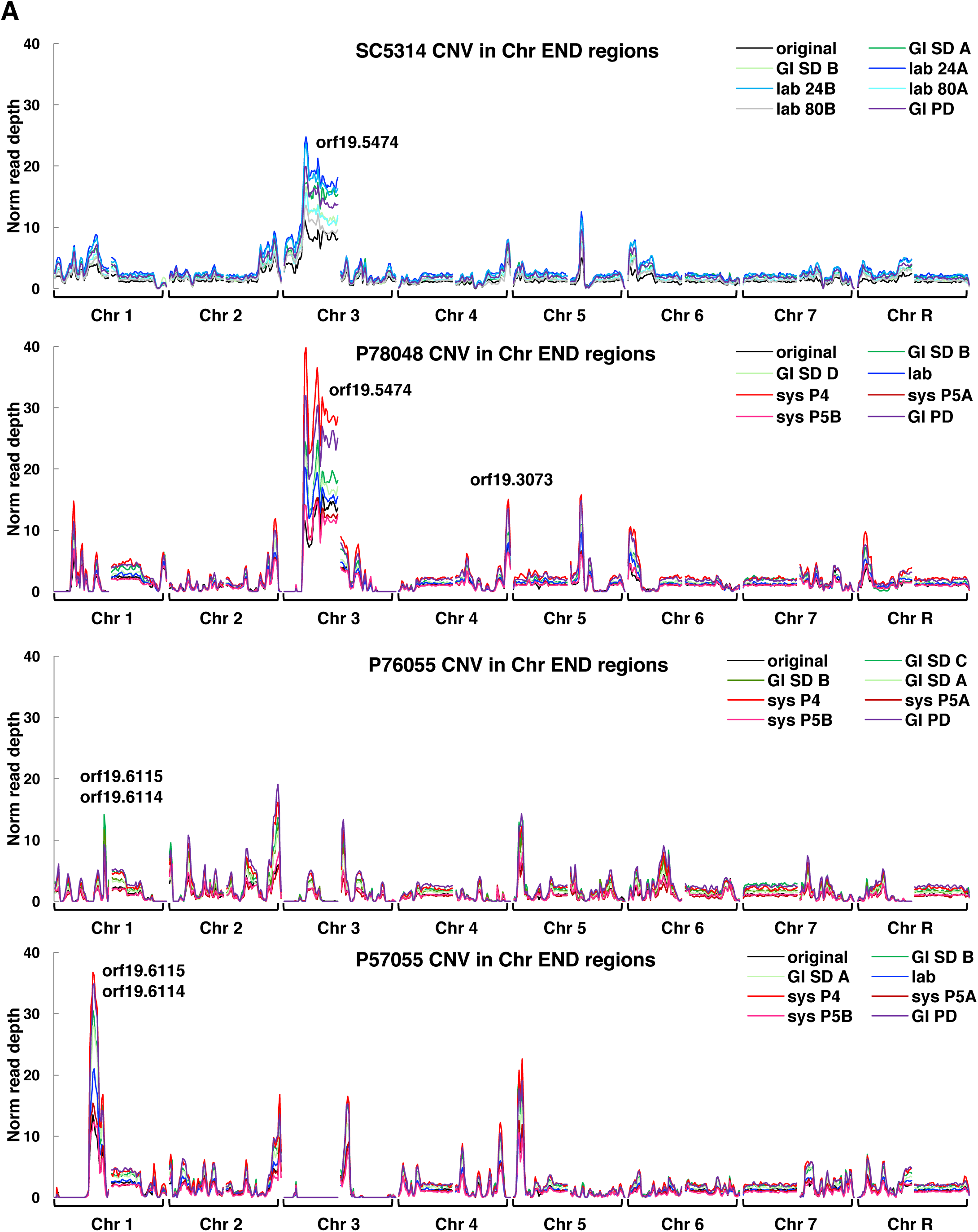
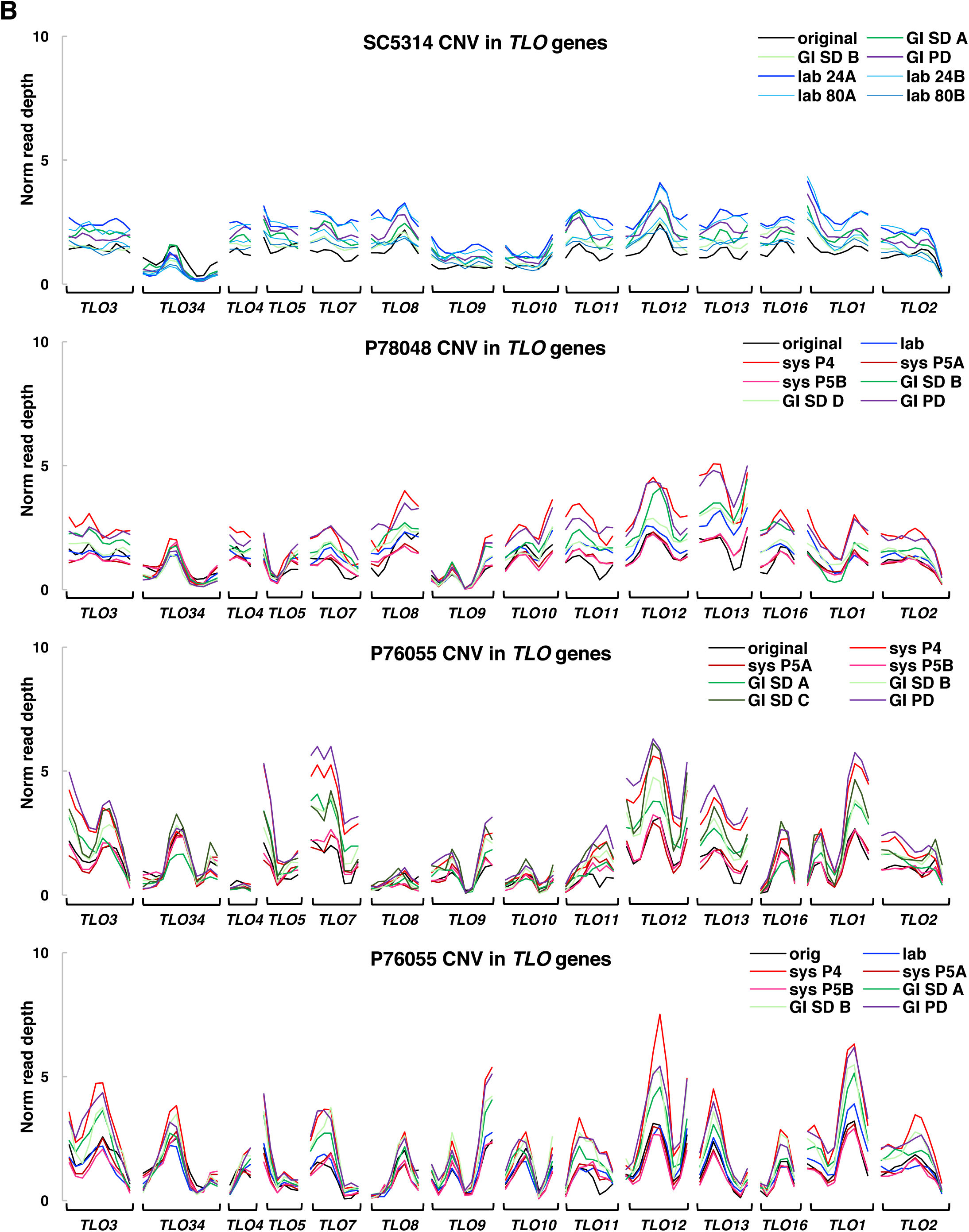
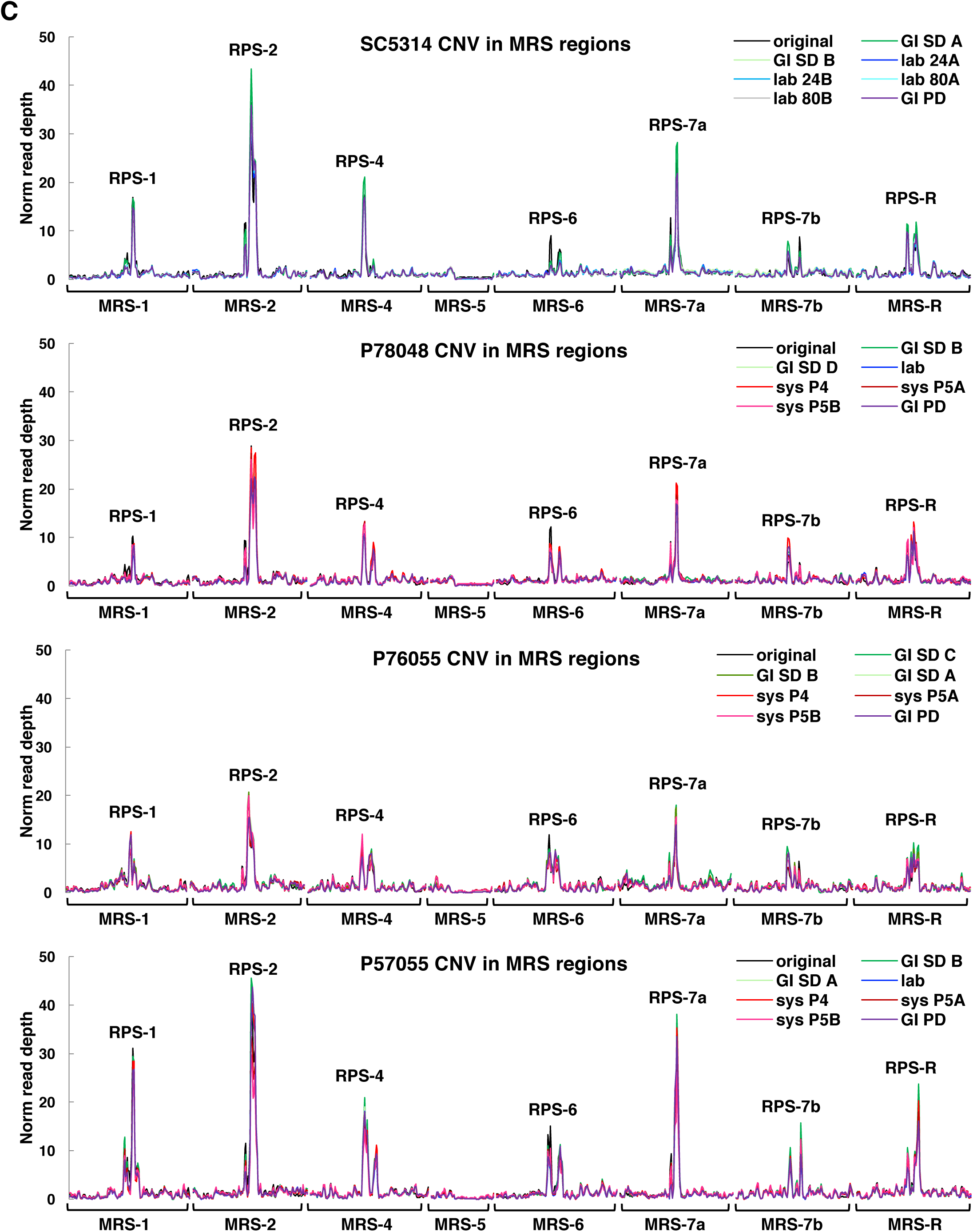
Copy number variation (CNV) patterns in specific genomic regions. Normalized read depths are shown for Chr END regions (terminal 5 kb at the ends of chromosomes, A), *TLO* genes (B) and MRS regions (C). CNV was determined by calculating the normalized read depth per 100 bp window. Copy numbers for original (starting) strains are shown in black lines.

## Supplementary Table legends

**Supplementary Table 1.** Strains used in microevolution experiments.

**Supplementary Table 2.** Sequencing and coverage information, including the frequency of heterozygous sites in each isolate. Sequencing variants were identified relative to the SC5314 genome reference strain.

**Supplementary Table 3.** Doubling times of the four lineages and *in vitro* calculation of generation times. Calculation of estimated *in vivo* generation times were based on (31) and (46).

**Supplementary Table 4.** Primers and results of the KASP genotyping assays. Probability values resulting from testing a binomial distribution model on the different types of mutations is also included.

**Supplementary Table 5.** List of nonsense and readthrough mutations identified during microevolution experiments.

**Supplementary Table 6.** Genomic coordinates for centromeres, chromosomal end regions, *TLO* genes, and *ALS* and GPI-liked genes, according to the *Candida* Genome Database (CGD).

**Supplementary Table 7.** Genomic coordinates for major repeat sequences (MRS), long terminal repeats (LTR) and genes within repeat regions according to the CGD.

**Supplementary Table 8.** Heterozygous (HET) and homozygous (HOM) regions of the four starting strains, as defined in (24). Included are also the overall genome heterozygosity levels based on these maps for the four strains.

**Supplementary Table 9.** Genes with known or predicted roles in DNA maintenance and DNA repair, including their genomic coordinates and CGD annotation.

**Supplementary Table 10.** Nonsense and stop codon mutations in DNA repair genes identified in the four starting isolates.

**Supplementary Table 11.** Mutations in DNA repair genes identified across microevolved isolates. Mutations involving LOH SNPs associated with LLOH events and resulting in nonsynonymous changes to the protein sequence are highlighted in red.

